# Lattice-Boltzmann Modelling for Inertial Particle Microfluidics Applications — A Tutorial Review

**DOI:** 10.1101/2023.04.10.536205

**Authors:** Benjamin Owen, Konstantinos Kechagidis, Sajad Razavi Bazaz, Romain Enjalbert, Erich Essmann, Calum Mallorie, Fatemehsadat Mirghaderi, Christian Schaaf, Krishnaveni Thota, Rohan Vernekar, Qi Zhou, Majid Ebrahimi Warkiani, Holger Stark, Timm Krüger

## Abstract

Inertial particle microfluidics (IPMF) is an emerging technology for the manipulation and separation of microparticles and biological cells. Since the flow physics of IPMF is complex and experimental studies are often time-consuming or costly, computer simulations can offer complementary insights. In this tutorial review, we provide a guide for researchers who are exploring the potential of the lattice-Boltzmann (LB) method for simulating IPMF applications. We first review the existing literature to establish the state of the art of LB-based IPMF modelling. After summarising the physics of IPMF, we then present related methods used in LB models for IPMF and show several case studies of LB simulations for a range of IPMF scenarios. Finally, we conclude with an outlook and several proposed research directions.

## 1 Introduction

Microfluidics involves the manipulation of small amounts of fluids in channels with dimensions between tens and hundreds of micrometres [1]. The precise handling of fluids and cells, the portability of devices, and the reduction or elimination of cross-contamination are some of the advantages of such miniaturised systems, making them appealing for lab-on-chip applications [2–4]. Over the past decades, several microfluidic methods have been developed — such as dielectrophoresis [5], magnetophoresis [6], acoustophoresis [7], thermophoresis [8], pinched flow fractionation [9], deterministic lateral displacement [10] and inertial microfluidics [11] — and applied to cell separation [12, 13], tissue engineering [14, 15], drug and gene delivery systems [16, 17] and clinical research [18–20].

While inertia is often negligible in microfluidic applications due to the small length scales involved, flow speeds in inertial microfluidics are significantly larger than their counterparts in non-inertial microfluidics [21]. As a consequence, the channel Reynolds number is typically of the order of several 10s or 100s, and a range of inertial effects can be exploited to manipulate the fluid, the suspended particles or both.

In inertial particle microfluidics (IPMF), which we focus on in this tutorial review, the aim is to manipulate suspended particles through inertial lift and drag forces. The most important inertial forces are i) the wall repulsion force, pushing particles away from nearby walls, ii) shear-gradient lift forces, usually pushing particles to regions of higher shear, and iii) drag forces in secondary flow fields that are caused by curved streamlines (*e.g.*, due to curved channels or obstacles in the flow) [11, 22–25]. Fig. 1 shows how, using these forces, particles can be focussed at off-centre lateral equilibrium positions, a phenomenon called the Segré-Silberberg effect [26, 27]. Particles can also be axially ordered into particle trains which have consistent inter-particle spacing. These inertial forces strongly depend on the properties of the particles and the shape of the underlying flow field, which in turn is governed by the channel geometry. Hence, channel design lies at the heart of many IPMF research efforts [25, 28, 29].

**Figure 1:**
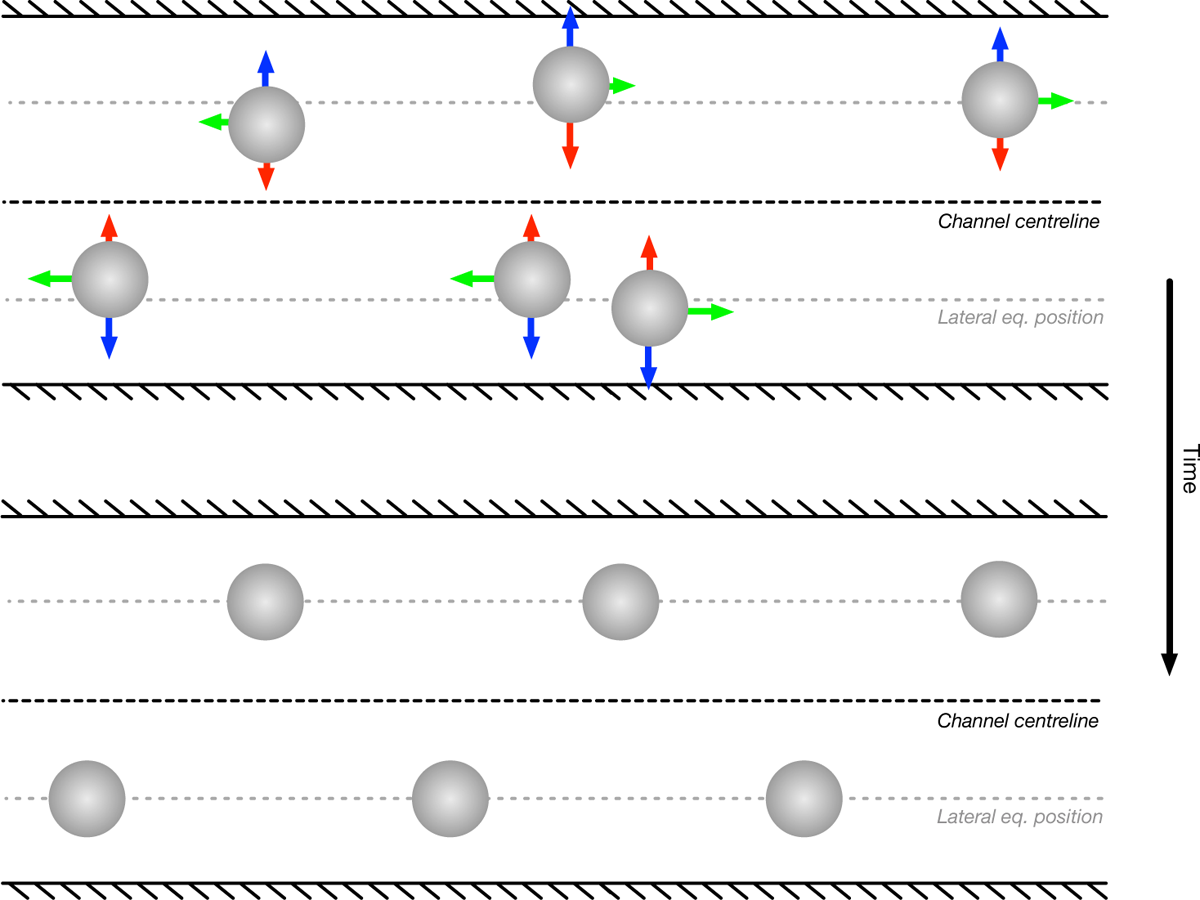
Visualisation of inertial forces acting on suspended particles. (Top) particles out of equilibrium and resulting forces at earlier times and (bottom) particles in equilibrium at later times. Each particle experiences a wall repulsion force (red) and a shear-gradient lift force (blue) which, once they are balanced, result in stable off-centre lateral equilibrium positions (Segré-Silberberg effect). Particles also interact hydrodynamically through their flow field distortions; the resulting drag forces (green) can lead to the axial arrangement of particles, in this specific case, the emergence of a staggered train with regular axial spacing. Once a stable train has formed, all inertial forces are balanced.

IPMF devices, for instance the one shown in Fig. 2, have been employed in a wide range of industries, such as microbiology [30–33], biochemistry [34–37], and biotechnology [38–42]. Many of these applications are devoted to the separation of a solid phase (cells, bacteria, and other particles) from a carrier fluid. For example, researchers have successfully demonstrated the isolation of circulating tumour cells [43–46], malaria parasites [47, 48], bacteria [49], and circulating fetal cells [50–52]. IPMF has been used for water filtration [53], dewatering of microalgae suspensions [54, 55], blood plasma separation [56–58], exosome sorting [59, 60], blood cell fractionation [61, 62], stem cell purification [63, 64], and the concentration of mammalian cells [65]. IPMF has also proven its value in flow cytometry applications and as a particle spacer [66–68], turning disordered dilute suspensions into orderly spaced particle trains that can be used for downstream processes [69–72], such as cell encapsulation in droplets [73, 74].

**Figure 2:**
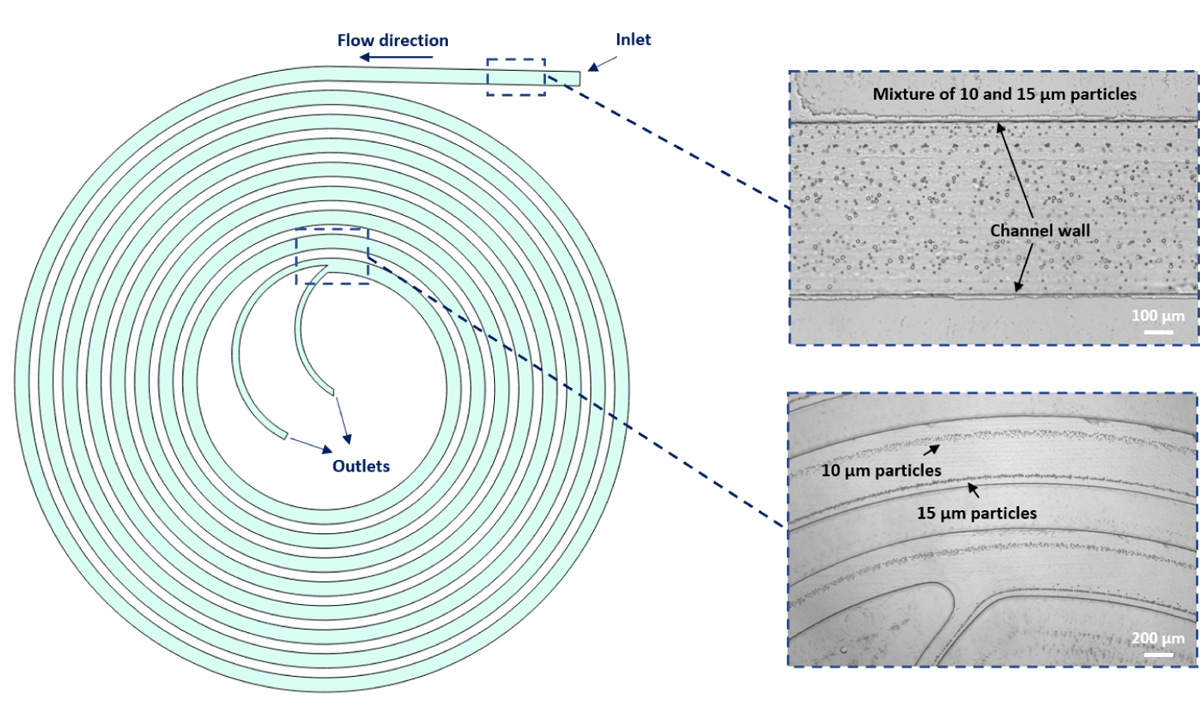
Suspension of particles in a spiral device. Disordered particles near the inlet. Separation of two types of particles near the outlet at a Reynolds number of 80. Different particle types have been focused in two different streams, allowing the separation of both populations. (Author provided) attack these challenges from two directions: i) simulate simple IPMF problems in detail to understand the underlying physics better and ii) develop reliable reduced-order models that are suitable for simulations with lower resolution. In this tutorial review we will focus on the former approach, which indirectly contributes to the success of the latter.

The governing equations of IPMF are non-linear, and there are no suitable analytical approaches to predict the dynamics of finite-size particles in realistic IPMF geometries [23, 75–77]. Due to the intricate relationship between device geometry, flow field and particle dynamics, most experimental IPMF research relies on time-consuming and costly trial-and-error approaches. The few existing design rules for IPMF devices have been established using particular device geometries; the range of applicability of these rules to more general designs is not well-understood [25, 68, 78–80]. Thus, in order to tailor geometries to particular needs (*e.g.*, the separation of specific tumour cells or bacteria from whole blood) and to optimise devices (*e.g.*, to reduce clogging or decrease the pressure needed to pump the fluid), computer simulations can be a powerful addition to the existing experimental expertise in the community.

Simulating IPMF systems comes with its own challenges. First, all IPMF scenarios are fluid-structure interaction (FSI) problems that require a high resolution of the flow field around the moving and possibly deforming particles in order to calculate the lift forces to a sufficient level of accuracy [81]. Second, most devices used today have channel lengths that are around three orders of magnitude larger than the particle diameters, and the assumption of periodic boundary conditions is often not appropriate for IPMF applications. Third, IPMF devices often have relatively small confinement (particle diameter divided by channel height) to avoid clogging [11, 82]. Thus, there are several relevant length scales that cannot all be resolved at the same time. It seems attractive to

Over the past 30 years, the lattice-Boltzmann (LB) method has emerged as an alternative to conventional computational fluid dynamics (CFD) approaches [83–87]. It has since been accepted that the LB method is a Navier-Stokes solver in its own right, and the method shows a number of advantages that makes it attractive for IPMF [88]. While conventional CFD methods directly solve the Navier-Stokes equations, the LB method is rooted in kinetic theory [89–93]. No pressure Poisson equation needs to be solved, which results in high numerical efficiency but also artificial compressibility [94, 95]. The local and kinetic nature of the LB method makes it suitable for problems with complex and moving geometries [96–99]. Due to its intrinsic properties, the LB method is particularly useful for fluid dynamics problems with Reynolds numbers between around 1 and several 100, which includes the typical range of Reynolds numbers found in IPMF. Unlike other particle-based counterparts, such as multi-particle collision dynamics and dissipative particle dynamics, the original LB algorithm is fully deterministic without thermal fluctuations, therefore advantageous for IPMF with its high Péclet number [100].

The scope of this tutorial review is to provide a comprehensive, yet concise overview of LB modelling for IPMF. We review the published literature (Section 2), define the physical and mathematical models underpinning IPMF (Section 3), describe the numerical methods (LB method in Section 4, particle methods in Section 5, FSI in Section 6, and additional numerical modelling considerations in Section 7), and provide four example cases that can be used for practice and validation purposes (Section 8). Finally, we provide an outlook with challenges and opportunities (Section 9). We assume that readers have a basic understanding of IPMF and the LB method. For those readers who have not, we provide relevant references throughout the text.

## 2 Overview of existing works

In this section, we focus on previous works in the field of inertial particle microfluidics (IPMF) using the lattice-Boltzmann (LB) method. We pay particular attention to the underlying physical observations and mechanisms, while the physical model and the numerical methods are detailed in later sections. The dynamics of particles in inertial microfluidics is largely affected by four main physical ingredients: the channel geometry, the degree of inertia (Reynolds number), the particle properties, and the concentration of particles. To maximise benefit to the reader and to avoid repetition, we first cover different types of geometries in Section 2.1 and then the role of particle concentration in Section 2.2. The effects of Reynolds number and particle properties (*e.g.*, size, softness, density) are included throughout both sections. Fig. 3 shows a timeline of important works that contributed to the development of LB-based IPMF modelling. Table A1 lists all works we identified that use LB to simulate at least one particle in a microfluidic channel at appreciable inertia.

**Figure 3:**
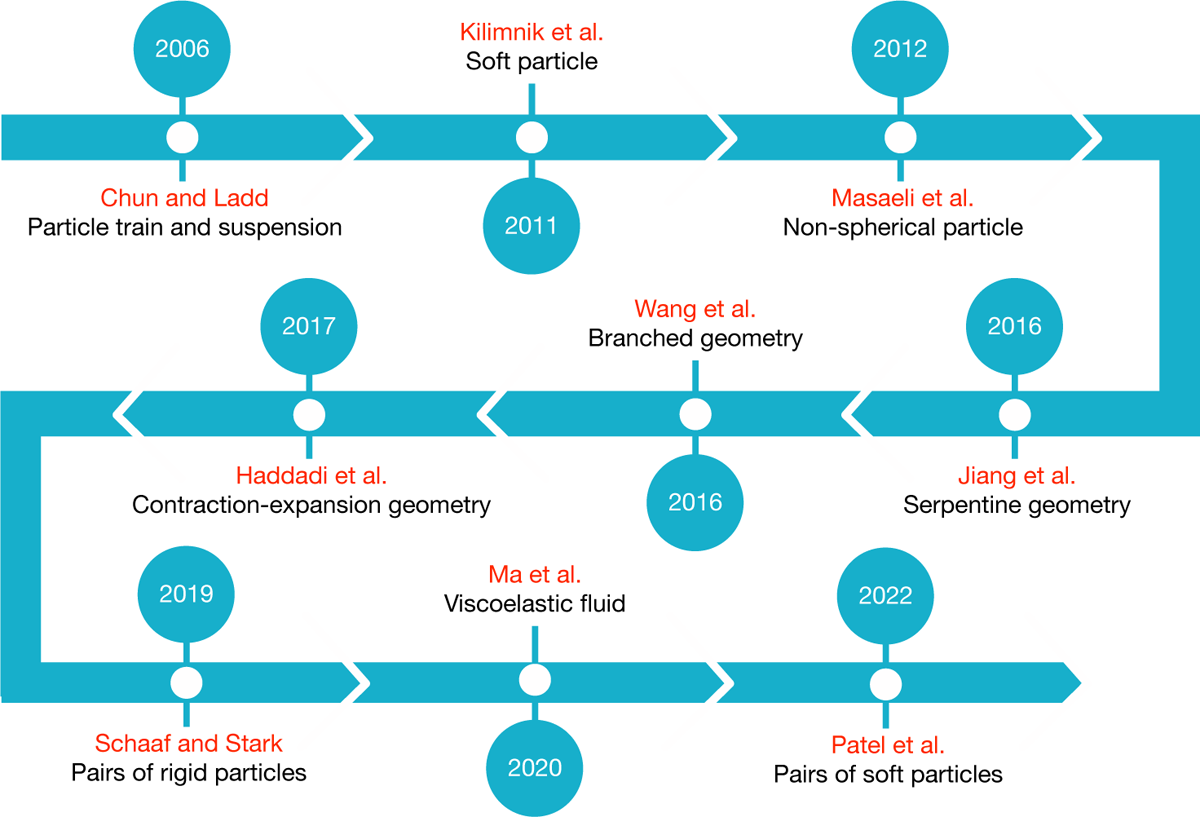
The timeline of important milestones in inertial particle microfluidics using LB. References to the relevant published works are as follows: Chun and Ladd [101], Kilimnik *et al.* [102], Masaeli *et al.* [70], Jiang *et al.* [103], Wang *et al.* [104], Haddadi *et al.* [105], Schaaf and Stark [106], Ma *et al.* [107] and Patel *et al.* [108].

A particularly important consideration is the dimensionality of the simulations. While all IPMF applications are intrinsically 3D, many authors use 2D simulations. Hydrodynamic interactions, which are crucial for suspension dynamics, are different in 2D and 3D. Hence, 3D simulations are essential for quantitative predictions. 2D simulations, however, can still shed light on fundamental mechanisms and inform further 3D investigations. Unless mentioned otherwise, all studies in this section are 3D.

**Figure 4:**
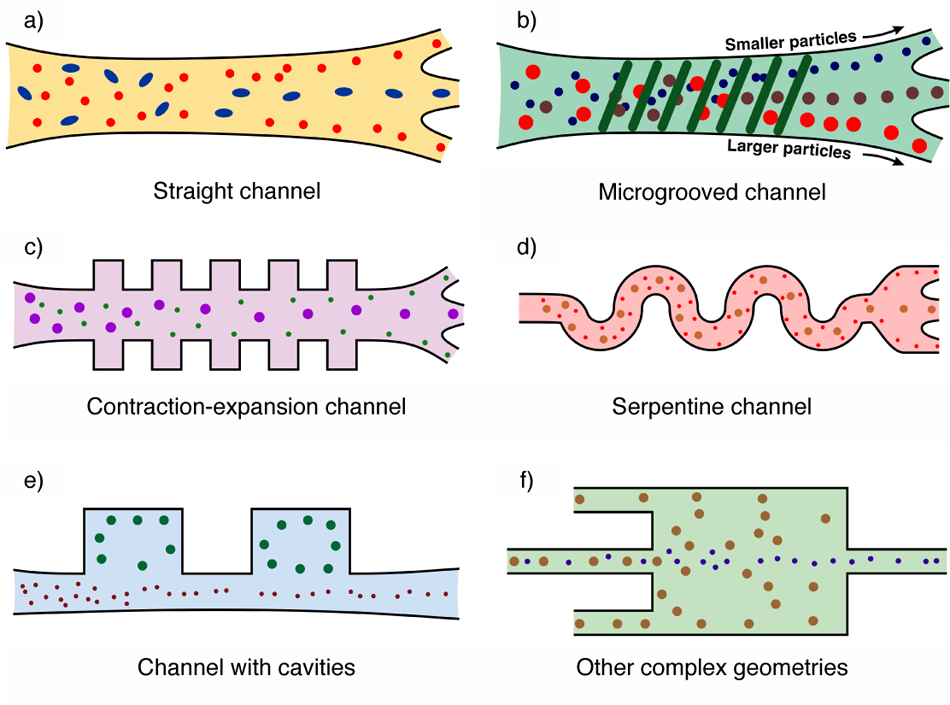
Commonly employed channel structures in inertial microfluidics: a) Straight channel, b) Microgrooved channel, c) Contraction-expansion channel, d) Serpentine channel, e) Channel with cavities, f) Other complex geometry. The flow is from left to right in all cases.

### 2.1 Microfluidic geometries

A key aim of inertial microfluidic devices is the manipulation of the dynamics and trajectories of suspended particles. Since particle dynamics is strongly affected by the flow field, which in turn is shaped by the device geometry, a wide range of geometries have been proposed (see Fig. 4). Factors such as the efficiency of particle focusing and separation, ease of fabrication and device footprint play important roles in choosing a suitable geometry. An open challenge is the design of optimised geometries for specific purposes, *e.g.*, the recovery of circulating tumour cells from whole blood. In the following, we provide a panoramic review of LB-based inertial microfluidic applications in a variety of channel types: straight channels (Section 2.1.1), straight channels with additional features (Section 2.1.2), curved channels (Section 2.1.3) and other geometries (Section 2.1.4).

#### 2.1.1 Straight channels with smooth surface

In order to decouple fundamental effects caused by the geometry and the particle behaviour in IPMF, it is advantageous to start with the simplest possible geometry: straight channels. Most channels used in 3D LB-based IPMF have a rectangular cross-section (referred to as ducts), reflecting their ease of fabrication compared to non-rectangular cross-sections. While other cross-sectional shapes have been explored in experiments to induce different particle lift force profiles, these ideas have not yet been picked up by the LB community [109, 110]. In the following, we highlight a few key works using straight channels.

Chun and Ladd [101] conducted the first LB-based study of IPMF, investigating the equilibrium positions of rigid particles within a square duct, both for single particles and dilute suspensions. The authors demonstrated that the equilibrium positions of single particles generally move closer to the channel centre with increasing Reynolds number, in agreement with theory and experiments [27]. It was also shown that the location of the equilibrium positions within the channel cross-section depends on the Reynolds number: particles favour the diagonal lines (corners), rather than the edges centrelines, at higher Reynolds numbers.

Prohm and Stark [111] investigated the effect of the aspect ratio of the channel cross-section in rectangular ducts. The authors observed that particles migrate to the middle of the longer edges when the aspect ratio decreases from unity to around 0.5, in agreement with previous experimental work [112]. This phenomenon effectively allows quasi-2D analyses for particle migration since particles were observed to stay on the midplane without external interactions. Later studies have taken advantage of the quasi-2D behaviour in order to reduce the number of independent parameters, thus facilitating the analysis of particle-particle interaction mechanisms in the inertial regime [106, 113].

Several studies investigated the effect of particle shape and density in straight channels. Hu *et al.* [114] demonstrated, in 2D, that elliptical and rectangular particles perform steady oscillations about their average lateral equilibrium position. Similar steady oscillations were also observed for non-neutrally buoyant particles at sufficiently large Stokes number [115, 116].

Straight channels have been used for the investigation of soft particles in inertial microfluidics for single particles [102, 117], pairs of particles [108, 113], particle trains [118], and particle suspensions [119]. A more detailed discussion of the effect of particle concentration is provided in Section 2.2.

There has been increasing interest in employing non-Newtonian fluids for IPMF. Straight channels have been used to demonstrate that the lateral equilibrium position of a particle is modified in a non-Newtonian fluid [114, 120]. A more detailed discussion of the effect of non-Newtonian fluids is provided in Section 2.2.

#### 2.1.2 Straight channels with geometric alterations or surface patterns

Channels with constrictions, cavities, grooves and similar features induce secondary flows, enabling more efficient manipulation of particles than in plain straight channels [121]. Here we summarise IPMF simulations of channels with additional features.

Mao and Alexeev [122] published the first work using LB for IPMF in a geometry more complex than a straight channel with smooth walls. The authors performed 3D simulations combining LB with the lattice spring model to investigate the motion of neutrally buoyant solid particles in a channel with diagonally aligned ridges (or grooves) on the channel walls. For Reynolds numbers up to 20, it was found that the ridges enhanced the separation of differently-sized particles.

3D simulations in contraction-expansion microchannels were performed to separate rigid particles of different sizes [123]. The smaller particles were affected by the secondary helical flow patterns and migrated closer to the walls. Contrarily, the larger particles stayed near the channel centreline. Jiang *et al.* [124] simulated a mixture of particles with two different sizes in 3D channels with different contraction-expansion ratios for Reynolds numbers between 15 and 120. It was found that particle focusing was improved at larger contraction-expansion ratios.

Haddadi and Di Carlo [105] explored the dynamics of a single or a dilute suspension of neutrally buoyant particles inside a cavity using 3D simulations. The Reynolds number was varied between 30 and 308, and the size of the vortex gradually increased with the Reynolds number. It was shown that the vortex is able to trap particles, especially larger ones. Jiang *et al.* [125] investigated a single rigid particle in a cavity through 3D simulations. The different observed particle entrapment modes result from an interplay of centrifugal and inertial lift forces.

More recently, Nizkaya *et al.* [126] considered particle migration in a straight channel with superhydrophobic striped walls in 3D. It was found that the superhydrophobic stripes change the lift forces acting on the particles and, therefore, their equilibrium positions.

#### 2.1.3 Curved or serpentine channels

Curved channels for IPMF have proven advantageous since the induced Dean flow contributes to the separation of particles and can reduce the required length for particle focusing by nearly an order of magnitude [24]. Curved geometries have also been shown to decrease the effective viscosity of the suspension [127]. However, simulating curved channels, in particular those with a large radius of curvature, is challenging, and only a limited number of studies using LB have been published to date. There is a strong need for simulating channels with large curvature radii in order to match experimental practice.

Jiang *et al.* [103] simulated suspension flow in a serpentine channel using the immersed-boundary method. In order to manage the complexity of the geometry, the authors simulated a single curved unit of the symmetric serpentine and applied periodic boundary conditions in flow-wise direction. Two populations of neutrally buoyant particles with different sizes were considered, and the Reynolds number varied between 25 and 100. At a low Reynolds number, the smaller particles were located closer to the sidewalls. With increasing Reynolds number and increasing Dean drag, the small particles were first focused closer to the channel centreline and then returned back to the side walls at a Reynolds number of 100. Larger particles remained near the channel centreline over the entire investigated range of Reynolds numbers demonstrating the dependence of particle size-based separation performance on Reynolds number.

Using 2D simulations in a serpentine channel, an empirical relationship was developed between the fluid/solid density ratio and the time taken for a particle to pass through the channel [128]. A critical value in the particle-to-fluid density ratio was found that allows a single rigid particle to traverse the channel faster. It was concluded that both the initial particle position and the value of the Reynolds number contribute significantly to the particle equilibrium position.

Ni *et al.* [129] simulated, and experimentally verified, the focussing of particles in an asymmetric serpentine channel with high aspect ratio. The authors observed that the periodic turn of the Dean flow causes the particles to migrate in waves within the channel and promotes the 3D single line focusing of the particles. It was concluded that 3D focusing of the particles near the centreline can be achieved irrespective of particle size while the focusing can be controlled by flow rate.

#### 2.1.4 Other types of geometries

There are hardly any limits to the diversity of geometries that can be used for IPMF applications. More complex geometries, such as those with multiple inlets and outlets, allow for more bespoke particle behaviour to be used in various applications.

Inspired from the shark’s skin, the entrapment of particles by the vortex in a riblet structure has been simulated in 2D at Reynolds numbers between 4.7 and 12 [130]. The authors found that flow pulsatility has a strong effect: particles were trapped in geometries with flat-edged walls in a steady flow, and smaller particles escaped the vortex under pulsating conditions. In the case of curved walls, particles remained trapped only at the lowest studied Reynolds number.

Wang *et al.* [131] investigated the motion of a hyperelastic capsule in a diverging T-shaped junction using the immersed-boundary method in 3D. The effects of capsule softness, Reynolds number and junction flow split ratio were considered. It was found that higher inertia causes the capsule to remain in the main branch, even when the side branch received a higher flow rate. Larger capsules had a higher probability of entering the side branch. Capsule softness introduced additional complexity; softer capsules show a stronger cross-streamline migration, making it possible to leave through the side branch under some circumstances.

Kechagidis *et al.* [132] studied the transient behaviour of a rigid particle passing through a cross-slot with two inlets and two outlets and a steady-state vortex located at the centre of the junction. The authors reported that larger particles and initial positions closer to the plane of vortex rotation lead to an increased residence time within the junction.

### 2.2 Particle concentration

While channel geometry determines the flow field and, therefore, the leading order contributions of lift and drag forces, hydrodynamic particle interactions are crucial and can lead to a variety of phenomena in IPMF. We first revisit the behaviour of single particles in more detail (Section 2.2.1), before reviewing particle pairs (Section 2.2.2), trains (Section 2.2.3), and suspensions (Section 2.2.4).

#### 2.2.1 Single particles

Single particle simulations are crucial in understanding the fundamental mechanisms behind IPMF. Removing all possible particle-particle interactions, it is possible to investigate the effect of parameters such as device geometry, particle properties (*e.g.*, shape, size, rigidity), and flow conditions.

##### Fundamental migration dynamics

Prohm and Stark [111] demonstrated that the eight equilibrium positions in a duct with a square cross-section previously identified by Chun and Ladd [101] are not all stable at the same time. In particular, it was found that small rigid particles migrate to face-centre equilibrium positions, while larger rigid particles migrate to diagonal equilibrium positions. Jebankumar *et al.* [115] incorporated particle density as an additional degree of freedom to the analysis. They demonstrated that, at a low Stokes number, non-neutrally buoyant particles behave in the same way as neutrally buoyant particles: these particles migrate to a steady lateral equilibrium position. However, at higher Stokes numbers, non-neutrally buoyant particles oscillate about a mean equilibrium position. Further work by Zhang *et al.* [116] showed that the oscillation amplitude increases with inertia. They observed that non-neutrally buoyant particles oscillate about the channel centreline if the oscillation is large enough for particles to cross the centre of the channel. Recent experimental work showed that, upon increasing the Reynolds number, the particle equilibrium positions first move closer to the channel walls before reversing the trend and moving back towards the channel centreline [133]. Yuan et al. [134] attributed this behaviour to the increasing size of the two vortices around the particle which, once large enough to get in contact with the wall, push the particle back towards the centre of the channel.

##### Effect of particle shape

Single particle simulations have been used to understand the effect of particle shape on inertial migration. Several works have investigated channel flows with non-circular particles in 2D [114, 135–138] and non-spherical particles in 3D [70, 139, 140]. Investigations of rigid biconcave objects have shown that their tumbling rotation leads to an oscillation of the lateral positions that depends on the particle size and aspect ratio [135]. Masaeli *et al.* [70] investigated the migration of ellipsoids of differing aspect ratios and demonstrated that the lateral equilibrium position depends on particle shape. Increasing the aspect ratio of ellipsoids has been shown to reduce the rotation frequency of the particle [137] and to facilitate the formation of stable trains (see Section 2.2.3) at high particle concentrations and for large particle size due to the high inclination angle of the ellipsoids in the train [141]. Recently, Nizkaya *et al.* [139] demonstrated that the equilibrium position of an oblate spheroid is shape-dependent with regard to its equatorial radius only. The authors also suggested a strategy to compare the behaviour of an oblate spheroid and a spherical particle. Li *et al.* [140] demonstrated that three different equilibrium behaviours exist for a single oblate spheroid: log-rolling, tumbling, and inclined log-rolling, with the latter disappearing with increasing Reynolds number.

##### Effect of particle deformation

Kilimnik *et al.* [102] were the first to model deformable particles using LB in IPMF, finding that particle equilibrium positions move closer to the channel centre when particles are softer. These observations were verified in 2D by Sun *et al.* [118]. Chen [142] expanded on this work by investigating the contributions of inertial and deformation-based migration, uncovering that particle migration is driven by competition between particle elastic contraction, fluid shear forces and fluid inertial stress. Schaaf *et al.* [117] investigated the lift profiles of particles across the channel width, showing that profiles for rigid and soft particles are similar. However, it was found that a significant difference between rigid and soft particles exists if an axial control force was introduced: rigid particles move towards the channel centre once they are slowed down, while soft particles do the opposite with the equilibrium position being independent of the degree of softness.

Apart from particle deformability, Takeishi *et al.* [143] further took the stress-free shape of soft particles into account, where biconcave red blood cells with various initial positions and orientation angles in a circular channel were modelled. The complex shape of the particle was found to introduce bi-stable motion regimes, depending on the particle initialisation, namely rolling and tumbling, the former of which impedes the inertial migration of the particle towards the wall whereas the later promotes such migration. Furthermore, the equilibrium position of the biconcave particle in the tumbling regime could be closely approximated by its spherical equivalent.

##### Effect of complex fluids

In recent years, there has been increasing interest in the inertial effects of particles in non-Newtonian liquids, such as shear-thinning [144–146] and shear-thickening liquids [147]. Focus has been placed on the reduced focusing length required by shear-thinning liquids [144], with particles suspended in these liquids migrating to equilibrium positions farther away from the channel centre [148]. Başağaoğlu *et al.* [136] have shown that particle migration is shape-dependent in non-Newtonian liquids. Investigations of stratified flows have demonstrated that the equilibrium position of the particle can be manipulated by varying the viscosity and flow rate of the two-component liquids [120]. Another method of manipulating the equilibrium position of particles is through the use of viscoelastic liquids. Ni and Jiang [149] demonstrated that the equilibrium position can be controlled through the elasticity number, the ratio between inertial and elastic forces, with increasing elastic number resulting in positions closer to the channel wall.

##### Behaviour in complex geometries

Single particle simulations have also been used to understand the effects of more complex geometries, such as ridged channels [122], bifurcating channels [131], channels with cavities [125], serpentine channels [128] and channels with superhydrophobic striped walls [126] (see Section 2.1 for details).

#### 2.2.2 Particle pairs

In addition to the Segré-Silberberg migration of a single particle, one observes that multiple particles align in the direction of the flow. When the particle number density increases, the particles form train-like structures with a characteristic axial spacing [68], see Fig. 5. These trains occur as a sequence of pairs where particles are located either on the same side (linear pairs) or on opposite sides of the channel (staggered pairs) [78]. Before discussing these trains in Section 2.2.3, we first need to establish the behaviour of particle pairs.

**Figure 5:**
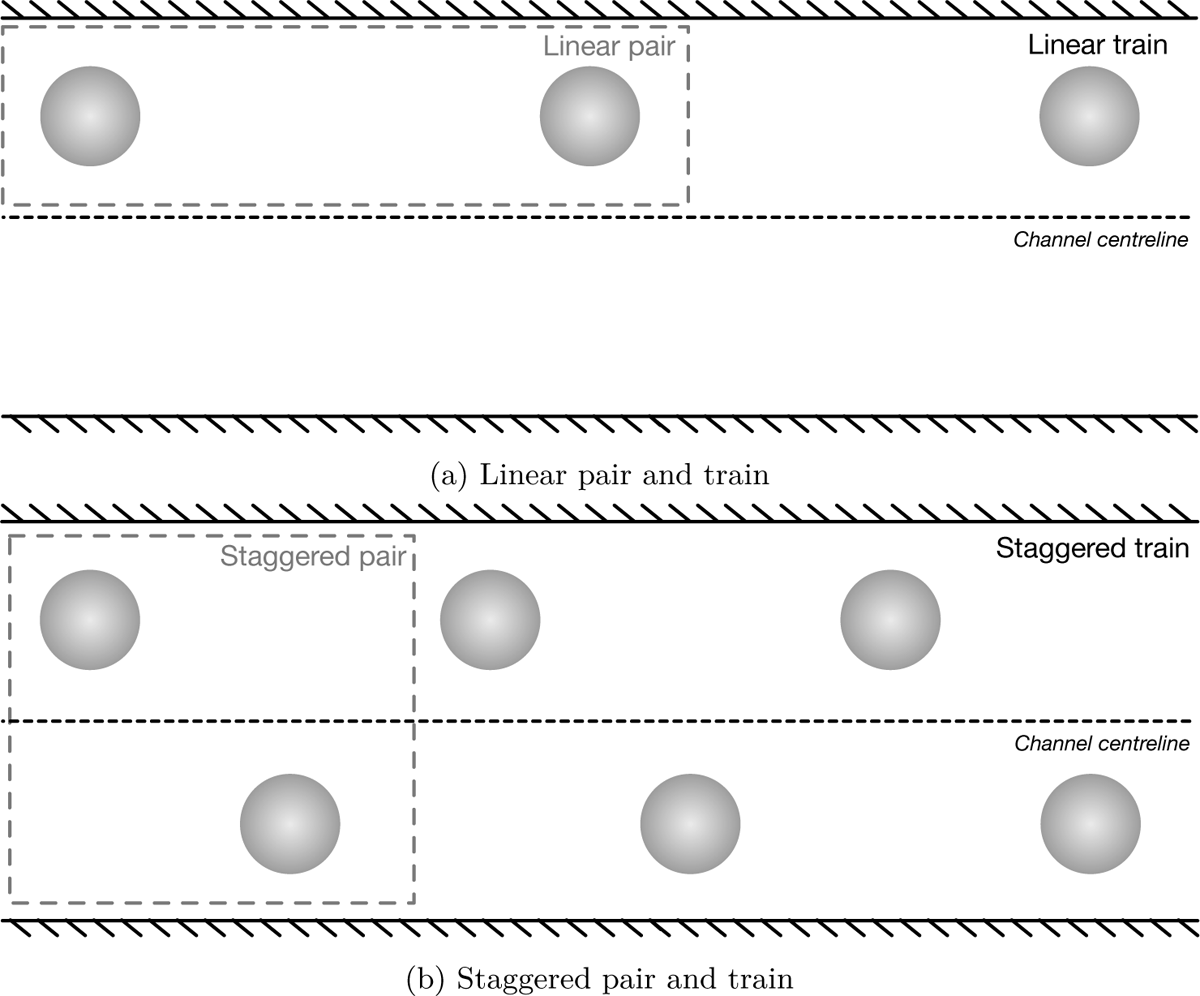
Illustration of particle pairs and trains. **a)** For a linear pair and train, particles are on the same side of the channel. b) For a staggered pair and train, particle positions alternate between both sides of the channel.

The analysis of particle pairs is typically done in rectangular channels since they are easy to fabricate and, unlike square channels, the number of equilibrium positions is reduced to two at the midpoints of the longer edges of the cross-section [111]. Particles tend to stay on the midplane once focussed there, thus simplifying the analysis. In such a configuration, staggered pairs assemble at a typical distance of around four particle radii [106, 113], while linear pairs have about twice that axial distance [106, 150].

The axial spacing of staggered pairs emerges as a combination of two effects: the inertial lift force pushes the particles towards their single-particle equilibrium position while the imposed channel flow and the viscous disturbance flow determine the axial distance [78]. Qualitatively, the formation of staggered pairs can be understood as one particle moving in the disturbance flow created by the other particle [151]. Humphry *et al.* [69] observed that a single particle creates a disturbance flow in its own frame of reference where an inwards spiralling eddy is formed around four radii upstream and downstream of the particle. The second particle moves along these streamlines while the inertial lift force drives the particle to the centre of this eddy [106, 152]. In such a stable configuration, both particles move with exactly the same axial velocity and the lift force is zero for both particles [106]. Without inertia, the particles perform undamped oscillations instead [153].

The axial distance of linear pairs is about twice that of staggered pairs. Though this behavior has been reported by multiple numerical and experimental studies [68, 78, 154], the stability of linear configurations is still unclear. Some LB simulations have shown that linear pairs are stable [68], which was explained by a minimisation of the kinetic energy of the fluid [152]. These results are also supported by recent experiments which were performed for Reynolds numbers between 1 and 4 [155]. However, a numerical analysis of the two-particle lift force profile reported no stable linear pair configuration, at least for the investigated particle-to-channel confinement [106]. Rather, when two particles are initialised on the same streamline at the equilibrium distance of a staggered pair, the particles increase their axial spacing until no longer interacting with each other [156]. This result agrees with early experiments [78] and recent 2D LB simulations [144] reporting a slow increase of the spacing for linear trains.

Recently, Patel and Stark [108] analysed how deformability influences the behavior of particle pairs. The authors observed that the presence of the second particle can change the stability of the single-particle equilibrium positions. Depending on the initial conditions, the leading particle may not move to an off-centre position, but rather migrates toward the channel centre. Li *et al.* [157] demonstrated that a deformable and a rigid particle are able to form a pair after a number of passing interactions, exploiting the numerical artifact of periodic images. Owen and Krüger [113] demonstrated that highly deformable particles form pairs for a greater range of initial positions than less deformable particles. However, the authors also showed that highly deformable pairs that migrate to the channel centreline do not attain stable axial distances and therefore cannot be considered stable.

Lin *et al.* [158] investigated pair formation of elliptical and rectangular particles. The authors found that increasing the aspect ratio of the particle moves the lateral equilibrium positions closer to the channel wall while also increasing the axial inter-particle spacing. Chen *et al.* [159] investigated pair formation in 2D for bi-disperse particles. They found that, for the values of Reynolds number and particle confinement studied, pairs do not form when the leading particle is smaller, whereas pairs can form when the leading particle is larger. Thota *et al.* [160] also investigated the formation and stability of pairs of particles of different sizes in 3D. It was found that pair formation of differently sized particles is determined by their initial lateral position and axial arrangement, while the stability and properties of the pairs depend on the particle size and their size ratio.

#### 2.2.3 Particle trains

With increasing particle line fraction — the proportion of the length, rather than the volume, of a channel segment covered by particles — pairs assemble into trains which form along the channel axis where the particles are located close to their single-particle equilibrium position [161, 162], see Fig. 5. These trains appear in staggered, linear, or mixed configurations. Similar to pairs, staggered trains have a characteristic axial spacing of about four particle radii between neighbouring particles while linear trains assemble at about twice that distance [68, 163].

For rigid particles, the train configuration seems to be largely independent of the Reynolds number. However, for deformable capsules, the Reynolds number has a strong influence on the train configuration as shown in recent 2D simulations [164]. When the Reynolds number is small, deformable particles migrate towards the channel centre and form linear trains. For higher Reynolds numbers, the equilibrium position shifts towards the walls and capsules form staggered pairs. Upon an increase of the line fraction the pairs move together and form staggered trains.

The formation of staggered trains depends on particle-to-channel confinement and Reynolds number, with larger confinement generally increasing the range of Reynolds number in which a staggered train can be stable [165]. When a staggered train with multiple particles is initialised with an axial distance larger than the equilibrium distance, it contracts via several mechanisms [156]. Initially, only the two leading and the two trailing particles, respectively, move together and slow down due to a collective drag reduction [166]. This effect causes the trailing pair to separate, and the leading pair moves closer to the next particle in line. The resulting three-particle cluster slows down further [156] as trains consisting of more particles move slightly more slowly than those with fewer particles. The trailing pair is eventually able to catch up with the leading particles, resulting in the final stable configuration.

Hu *et al.* [144] analysed the formation of staggered trains using 2D LB simulations and found that particles do not have a fixed axial distance but rather perform oscillations around their equilibrium position [144]. The amplitude of the oscillation increases with the Reynolds number. Liu *et al.* [161] identified two different distribution patterns in staggered trains depending on particle concentration, a continuous pattern where uniform spacing between all particles exists, and a discontinuous pattern where a larger spacing exists between two particles, effectively breaking the train.

While the axial spacing in linear trains is characteristic, it has been reported that the distance slowly increases with time [78, 144, 156]. Linear trains are not perfectly aligned with the channel axis but have a slight inclination where trailing particles are pushed closer to the walls compared to the single particle equilibrium position. In an unstable linear train, the leading particle experiences a lift towards the channel centre, resulting in a slightly higher speed and a slow increase in the axial distance [144, 156].

Recently, particles of different shapes have been shown to form linear trains in 2D [167] while the formation of linear trains consisting of differently sized particles has been studied in experiments and simulations [168, 169]. Both studies showed that the trains behave similarly to homogeneous trains as long as the ratio of the particle diameters is close to unity. When the ratio increases beyond two, the smaller particles form pairs or triplets in the gaps between the larger particles. The pairs or triplets of small particles keep their position within the trains while the individual particles oscillate about their common lateral equilibrium position [169]. Most bi-disperse trains have been observed to have a large leading and a small trailing particle [168]. For large size differences, only the larger particles form trains, and the inertial focusing of the smaller particles is inhibited by the presence of the larger ones.

#### 2.2.4 Particle suspensions

The primary objective in many applications of IPMF is to focus particles into trains in a high-throughput fashion (*cf.* Fig. 2). Since Chun and Ladd [101] observed that particles in dilute suspensions focus at symmetrically placed equilibrium positions in a duct flow, a number of studies have investigated the effect of particle concentration on train formation. Kahkeshani *et al.* [68] found that increased concentration causes the inter-particle spacing to decrease until train formation is no longer possible. Feng *et al.* [164] showed that the formation of trains in dilute suspensions depends on particle concentration, Reynolds number, and particle softness. Huang *et al.* [170] further demonstrated that, once the line fraction was too large for more particles to enter the train, additional particles were likely to locate close to the channel centre, moving at a larger axial velocity than particles in the train.

Focussing and separation characteristics of particle suspensions are often used to assess and optimise the performance of complex geometries in IPMF. Examples include microcavities [105], serpentine flows [103], contraction-expansion flows [124] and channel networks [171]. See Section 2.1 for more details on the effect of geometry on IPMF.

For non-dilute suspensions (*i.e.*, at concentrations typically above the threshold below which stable trains can exist) under inertial conditions, the lateral migration of the suspension is important for optimising high-throughput particle enrichment and separation. Krüger et al. [119] simulated a capsule suspension at 10% volume concentration. The authors demonstrated that particle deformation and inertial effects both cause lift forces that can compete with or complement each other, resulting in reduced off-centre particle focusing. Further work found that inertial migration decreases with particle concentration in dense suspensions (volume concentration between 5% and 50%) [172]. Inertial migration was also observed to decrease due to agglomeration of particles with adhesive properties [173] while the focusing length was observed to increase with Reynolds number [174]. Millet [175] determined that the multi-directional confinement of the suspension hinders inertial focusing due to the capsule-free layers that develop in the two transverse directions. The thickness of capsule-free layer in a given cross-section depends on the wall length (in a transverse direction). As a result, a non-square cross-section (i.e. rectangular) has a non-homogeneous capsule-free layer.

Particles in non-dilute suspensions can also be separated based on their individual properties. Sun *et al.* demonstrated that particles of different size [176] and deformability [177] in non-dilute suspensions migrate to distinct lateral equilibrium positions in straight channels and therefore can be separated. Particles of different shapes can also be separated through the same mechanism, with the shape-dependence of lateral equilibrium positions found to be larger when particles are suspended in a pseudo-plastic rather than a Newtonian fluid [136].

## 3 Physical model

Having summarised the variety and richness of emerging behaviour of IPMF problems in Section 2, we now turn our attention to the underlying physical mechanisms that need to be included in any model of IPMF with resolved particles. This section summarises the underlying assumptions, physical model, and governing equations for IPMF. We need to consider the fluid (Section 3.1), the particles (Section 3.2), and the boundary conditions (Section 3.3). As a general note, although all inertial microfluidic experiments are three-dimensional (3D), there exist several two-dimensional (2D) LB-based models of IPMF. While any realistic IPMF application requires 3D simulations, 2D simulations can be useful for the study of fundamental mechanisms, for instance the migration of particles in channel flow.

### 3.1 Fluid model

The fluid model normally comprises incompressible Newtonian fluids governed by the continuity and Navier-Stokes equations (Section 3.1.1). We also briefly comment on external forces (Section 3.1.2) and non-Newtonian liquids (Section 3.1.3).

#### 3.1.1 Governing equations

In IPMF, we can assume that the fluid is an incompressible viscous liquid in the continuum limit. Since the liquid can be considered isothermal with negligible viscous heating, the energy equation can be neglected, and only the mass and momentum balance equations are relevant [178]; they take the form of the incompressible continuity and Navier-Stokes equations, respectively:

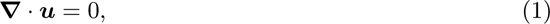

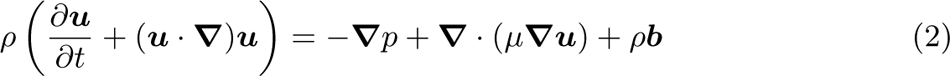

where *ρ* is the density, ***u*** is the velocity, ***b*** is an external body force density, *p* is the pressure, and *µ* is the dynamic viscosity (which is not necessarily constant). Section 4 describes the LB method as a numerical method to solve the Navier-Stokes equations.

We can define the channel Reynolds number as

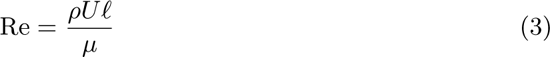

where *U* is a characteristic velocity (for example the average flow velocity in the channel) and *f* is a characteristic length scale, typically the smallest channel dimension. In IPMF applications, Re is usually in the range 10–500.

#### 3.1.2 External forces

IPMF in its original form is a passive method, *i.e.*, particles experience only fluid drag and lift and no other external forces. Several research groups have combined IPMF with active methods that involve electromagnetic or other forces to manipulate the flow and particles therein (see [179] for a review of active methods). In the following, we will not discuss forces related to active methods and instead focus on passive IPMF.

Gravity can usually be neglected in IPMF. The sedimentation speed *v* of a small spherical particle [180] (radius *a*, density *ρ*_p_) settling in a viscous fluid (viscosity *µ*, density *ρ*) can be estimated by equating the Stokes drag and buoyancy forces (gravity *g*): *v* = 2*a*^2^(*ρ*_p_ − *ρ*)*g/*(9*µ*). In a typical IPMF application, we expect the system to have dimensions of the same order as those in Table 1, resulting in a sedimentation speed of *v* ∼ 20 *µ*m*/s*. Typical particle advection speeds are ~ 1 m*/*s, making the effect of buoyancy orders of magnitude smaller than the effect of advection.

**Table 1:** Typical properties of liquids and particles in IPMF devices

|  |  |
| --- | --- |
| Particle radius, $a$ | $\sim 10 \mu\text{m}$ |
| Viscosity, $\mu$ | $\sim 1 \text{ mPa s}$ |
| Relative density, $\rho_p - \rho$ | $\sim 100 \text{ kg m}^{-3}$ |
| Gravity, $g$ | $\sim 10 \text{ m s}^{-2}$ |
| Flow velocity, $U$ | $\sim 1 \text{ m s}^{-1}$ |

#### 3.1.3 Non-Newtonian liquids

The vast majority of IPMF applications involve Newtonian liquids. However, several important biological fluids are non-Newtonian, at least over a limited range of shear rates, such as blood plasma [181, 182]. There has been a recent interest in combining inertial with non-Newtonian effects to modify particle focusing and separation behaviour [183–185]. See Section 4.4 for relevant LB-based papers.

### 3.2 Particle model

Certain particle properties are relevant for their dynamics in inertial flows (Section 3.2.1). Beyond these general properties, we distinguish between rigid particles (Section 3.2.2) and deformable particles (Section 3.2.3).

#### 3.2.1 General properties

Since the primary area of application of IPMF is the processing of biological cells, particle density is usually within 5–10% of the density of the suspending liquid. Typical particle radii *a* range from around one to 15 microns. Due to high flow speeds in IPMF, the Péclet number is large, and particles are non-Brownian. In cases where particle inertia is important, it is common to use the Stokes number, St, which is the ratio of the particle response time in the flow to a characteristic flow time scale (*e.g.*, advection) [115, 116].

A key parameter is the particle-to-channel confinement

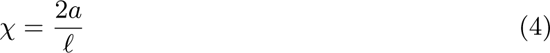

where *f* is a characteristic length of the channel cross-section. It is often convenient to define the particle Reynolds number Re_p_ which characterises the strength of inertia on the scale of the particle, rather than the channel. A common definition is Re_p_ = Re*χ*^2^, although various alternatives are used throughout the literature.

Particle concentration in rheology is normally given as a volume fraction *φ*, but in IPMF the line fraction *φ*_l_ is usually more important since it determines particle train formation which can happen even at small values of *φ*.

#### 3.2.2 Rigid particles

While most biological cells deform under the high stresses occurring in IPMF, rigid artificial particles are often used to characterise the focusing and separation characteristics of IPMF devices. In the majority of cases, rigid particles are spherical and fully characterised by their radius *a* and density *ρ*_p_. Non-spherical rigid particles can play an important role in IPMF, *e.g.*, ellipsoidal particles or hardened red blood cells, see Section 2.2.1.

The dynamics of rigid particles is determined by Newton’s equations of motion and rigid body dynamics (in the stationary reference frame):

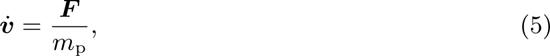

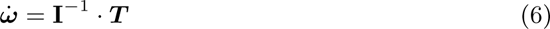

where ***v*** is the velocity, *m*_p_ is the mass, ***ω*** is the angular velocity and **I** is the inertia tensor of the particle. The total force and torque acting on the particle are denoted by ***F*** and ***T***, respectively. Since buoyancy is negligible and electromagnetic effects are usually absent, the forces and torques acting on a particle merely arise from fluid stresses (see Section 3.3).

#### 3.2.3 Deformable particles

Nearly all biological particles used in IPMF are deformed under high fluid stresses. Since deformable particles in fluid flow behave substantially differently than rigid particles, appropriate models for deformable particles must be considered in certain scenarios.

##### Types of deformable particles

One key application of IPMF is the processing, focussing and separation of biological cells, such as white blood cells (WBCs), circulating tumour cells (CTCs) and red blood cells (RBCs) [186]. Compared to WBCs and CTCs, RBCs have much simpler mechanical properties, which led to a number of accurate numerical RBC models (Section 5.3). RBCs are formed by a compound membrane comprising a lipid bilayer and a supporting cytoskeleton, while the interior consists of a viscous concentrated haemoglobin solution [187]. Modelling WBCs and CTCs as deforming particles accurately, however, is an ongoing challenge since these cell types have internal structures, more complex shapes and richer dynamics [104, 188].

Instead, the research community is often focussing on simplified models for soft particles. For example, capsules are hyperelastic membranes enclosing a liquid that may be different from the suspending liquid. Vesicles are lipid mono- or bilayers enclosing a liquid. Unlike capsule membranes, vesicle membranes are viscous and incompressible. Both capsules and vesicles have been used as models for RBCs due to similar properties and dynamic behaviour [189, 190]. For a comprehensive summary of the properties of capsules, vesicles and RBCs we refer to [189, 191–194].

##### Governing physical effects

The dynamics of deformable particles is governed by a number of physical effects. Biological membranes, like those of the RBC, are usually viscoelastic, incompressible and show a finite resistance to bending [187, 195, 196]. The corresponding material parameters are *κ*_s_ (elastic shear resistance of the membrane), *κ*_b_ (bending modulus) and *η*_m_ (membrane viscosity). The capillary number is the ratio of fluid stress in shear flow (shear rate *γ*) to membrane elastic stress:

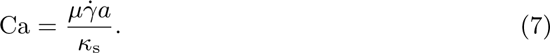

For RBCs (with radius *a* = 4 *µ*m), the ratio of elastic and bending moduli obeys *κ*_s_*a*^2^*/κ*_b_ ≈ 400 [192, 197, 198]. The internal dynamics of red blood cells, which are filled with a haemoglobin solution rather than carrying a nucleus, is determined by the cytoplasmic viscosity *η*_in_ that is about 5–7 times higher than that of water or blood plasma [199, 200]. The viscosity contrast is defined as Λ = *η*_in_*/µ*.

Other biological cells, such as leukocytes, have an internal structure with a nucleus (eukaryotic cells), organelles and microtubules. Nearly all deformable particles in IPMF do not change their volume in flow since they are filled with an incompressible medium and the membranes are impermeable to water on the time scales relevant to flow in IPMF devices. While vesicles, which are made of a liquid incompressible membrane, have a constant surface area, capsules can undergo surface stretching which is characterised by an elastic dilation modulus *κ_α_*.

##### Red blood cell membrane model

In the following, we will present physical models of the RBC membrane. Various models for the RBC have been developed over the past decades, some of which are generic enough for modelling other types of cells that can be considered as capsules or vesicles. Detailed models of leukocytes in IPMF have not yet been proposed.

Existing RBC models can be classified into two categories: continuum-level models and spectrin-level models. The continuum-level models are constructed from constitutive laws that describe the cell membrane as a thin and elastic shell separating the cytoplasm and the suspending medium. Common models in this category include the Skalak model [201] and the neo-Hookean model (a case of the Mooney–Rivlin model under small deformations) [202, 203]. The spectrin-level models, as the name suggests, mimic the spectrin-link network of the cytoskeleton supporting the lipid bilayer in the membrane [204, 205]. In spectrin-level models, the membrane is represented by a mass-spring system, which often needs to be coarse-grained to constrain the otherwise prohibitive computational cost [206].

We focus on the continuum-level model in more depth since the spectrin-level model has not been used for LB-based IPMF studies. The corresponding numerical model will be discussed in Section 5.3.1. Starting from the undeformed shape of the RBC, any deformation of a two-dimensional membrane element can be quantified by the two principal stretch ratios *λ*_1_ and *λ*_2_.

Assuming that the elastic properties of the RBC membrane are isotropic, each membrane element has only two physically relevant parameters (shear and dilation) which are often written as the strain invariants *I*_1_ = *λ*^2^_1_ + *λ*^2^_2_ − 2 and *I*_2_ = *λ*^2^_1_*λ*^2^_2_ *−* 1. In the following, we assume that the RBC membrane is hyperelastic [207]; see [208–210] for viscoelastic models of the RBC membrane. Any shear or bending deformation of the RBC is associated with an increase of the free energy of the RBC membrane: *E* = *E*_S_ + *E*_B_ where *E*_S_ and *E*_B_ are the strain and bending contributions, respectively.

Skalak’s model [201] is the most popular strain energy model for RBCs:

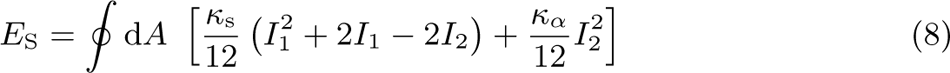

where the integration is performed over the closed RBC surface. The bending energy is often approximated by Helfrich’s model [211]:

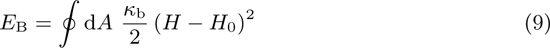

where *H* is the trace of the surface curvature tensor and *H*_0_ is the spontaneous curvature, a local property of the membrane. Since the total RBC surface area and volume remain nearly constant, constraints on the total cell volume and surface area are usually added in the form of stiff harmonic potentials [212]. Finally, the forces acting on each element of the RBC surface can be calculated through the principle of virtual work [213, 214].

### 3.3 Boundary conditions and fluid-structure interaction

Chemical transport and diffusion are often not part of IPMF applications. Thus, we focus on hydrodynamic boundary conditions only. There are generally three different types of boundaries that need to be considered in IPMF applications: 1) the boundary condition at the surface of the device; 2) the boundary condition at the surface of the moving and possibly deforming particles; 3) inlet and outlet conditions since IPMF devices are open systems, as illustrated in Fig. 6.

**Figure 6:**
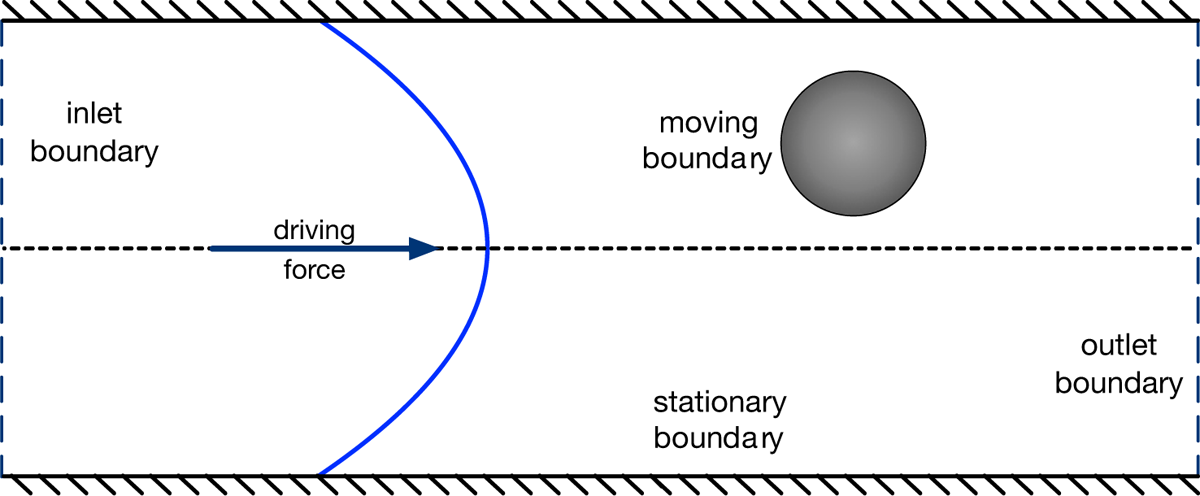
Different types of boundaries that need to be considered in IPMF, here illustrated for a straight channel: 1) the boundary condition at the stationary surface of the device; 2) the boundary condition at the surface of the moving and possibly deforming particles; 3) inlet and outlet conditions which may coincide with each other if periodic conditions are used. The flow (illustrated by the curved blue line) is usually driven by a pressure drop or driving force.

#### 3.3.1 Device-fluid boundaries

Device surfaces are normally assumed impermeable and satisfy the no-slip condition. The device surface can be considered rigid and immobile in most cases, hence the interaction of the device and the flow is fully characterised by the stationary surface and the no-slip condition. See Section 6.1.1 for the numerical treatment of the boundary condition at the surface of the device.

#### 3.3.2 Particle-fluid boundaries

The motion and deformation of particles both affect the flow and are affected by the flow, therefore defining an FSI problem. While the instantaneous particle surface shape imposes the no-slip condition, the particle translation, rotation and deformation are governed by hydrodynamic forces and torques. For rigid particles, these effects are described by (Eq. 5) and (Eq. 6). In IPMF applications, forces and torques acting on particles are usually of hydrodynamic origin, hence

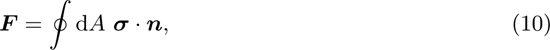

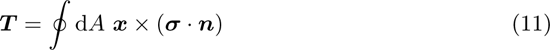

where ***σ*** is the fluid stress tensor with components *σ_αβ_* = − *pδ_αβ_*+ *µ*(*∂_α_u_β_* + *∂_β_u_α_*), ***n*** is the surface normal vector pointing into the surrounding fluid and ***x*** is the position of a point on the particle surface. The numerical treatment of the FSI problem is probably the most challenging aspect of IPMF modelling, see Section 6.2.

When particles come very close to other particles or the device surface, other forces might become significant, either for physical reasons (friction forces, van-der-Waals forces) or for numerical reasons (particle collision detection and overlap handling). Due to the typically low volume concentrations and large fluid stresses in IPMF, additional physical forces are normally not relevant. In simulations, however, additional lubrication or repulsion forces are often used to keep particles from overlapping, see Section 7.1.

It is possible to use under-resolved particles with appropriate drag and lift force models instead of particles with resolved FSI [215, 216]. Although under-resolved particle models are desirable for the simulation of large geometries that would otherwise be too expensive to be simulated, it is a major challenge to find suitable drag and lift force models that are accurate in general flow fields and in the presence of other particles. Since the study of the dynamics of resolved particles can be used to construct effective drag and lift force models, it is currently indispensable to focus the community’s modelling efforts on resolved particles.

#### 3.3.3 Inlet and outlet boundaries

Finally, the inlet and outlet conditions play an important role in IPMF (see Section 6.1.2 for their numerical treatment). In the vast majority of realistic modelling scenarios, only subsets of a device are of interest or can be afforded in simulations. Therefore, the flow on the inlet and outlet planes of the chosen subset must be specified. If the subset consists of a straight channel or is a unit cell of a channel with periodic features (*e.g.*, a serpentine channel), periodic boundary conditions are normally the most suitable and straightforward choice. Using periodic boundary conditions, any fluid or particle leaving the numerical domain on one side enters on the other side. Physically, the simulated system is an infinite array of unit cells where the unit cell is defined by the actual simulated domain.

If periodic boundary conditions are not appropriate, for example, if the device subset has a complex shape, it is necessary to impose velocity or pressure conditions at the inlet and outlet. Unless the flow field on the inlet plane is known, modellers usually need to impose the fully developed velocity profile for a given channel cross-section which assumes the absence of any upstream flow perturbations. Closed-form time-independent solutions to the Navier-Stokes equations for the duct pipe are known for many geometrically simple cross-sections [178, 217], and these solutions are the natural choice for a fully developed velocity profile. Since the downstream range of flow perturbations increases with Reynolds number [218, 219], this assumption can be inappropriate for IPMF applications, making the entire simulation invalid in the worst case. The treatment of the outlet is conceptually simpler since the upstream range of flow perturbations outside the simulated domain is small and the flow field on the outlet plane is largely determined by the flow inside the simulated domain. A common alternative outflow condition to periodicity is a zero-gradient condition [125, 131]. A further complication of non-periodic boundary conditions is the treatment of particles entering and leaving the subset (see Section 7.2 for numerical details).

## 4 Lattice-Boltzmann method for fluid flow

After having summarised the physical model of IPMF in Section 3, we will now outline the lattice-Boltzmann (LB) method for fluid flow. We focus on those LB aspects and features that are relevant for simulating IPMF: LB essentials (Section 4.1), collision operators (Section 4.2), forcing schemes (Section 4.3), and non-Newtonian fluids (Section 4.4). Numerical boundary conditions and FSI approaches are covered in Section 6 since they require a detailed separate discussion. We do not present a comprehensive LB summary here since there exist various suitable introductory texts, such as [88, 94, 220, 221]. Several open-source LB codes are available, some of which can be used for IPMF simulations, for example, OpenLB [222], Palabos [223] and waLBerla [224].

### 4.1 Lattice-Boltzmann essentials

The main aim of LB is to solve the Navier-Stokes equations governing fluid mechanics (Section 3.1) by discretising the Boltzmann equation in space, time and velocity space, and by replacing the Boltzmann collision operator by a simplified relaxation step. As a result, the probability distribution function *f* (***x, v****, t*) becomes a finite set of discrete populations *f_i_*(***x****, t*) that can move on a lattice with the corresponding discrete velocities ***c****_i_*. The stencil defined by the *d* spatial dimensions of the problem and the number *q* of discretised populations is called D*d*Q*q*. For example, a common discretisation in 3D space involves 19 populations at each lattice point, hence the associated stencil is called D3Q19 [225].

In the LB method, the populations *f_i_* propagate and collide on a regular lattice. The corresponding evolution equation, the LB equation, is generally written as

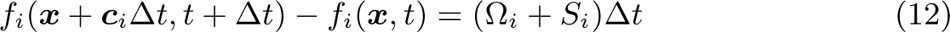

or

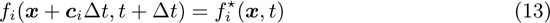

where Δ*t* is the time step, Ω*_i_* is the collision operator, *S_i_* includes any source terms, and the *f\**(***x****, t*) = *f_i_*(***x****, t*) + (Ω*_i_* + *S_i_*)Δ*t* are called the post-collision populations. The collision operator in the LB method is usually modelled as a relaxation process in which the populations relax towards a local equilibrium state *f* ^eq^. Details about the collision operator and the equilibrium distribution are summarised in Section 4.2. The source term includes any external forces, such as gravity, but also those forces that come from FSI schemes, such as the immersed boundary method (Section 6.2.2). The inclusion of forces in the LB method is discussed in Section 4.3. The left-hand-side of (Eq. 12) is called the propagation or streaming step as it describes how a population *f_i_* moves from one point ***x*** to its neighbour by the corresponding distance ***c****_i_*Δ*t* during a time step Δ*t*.

The macroscopic variables of fluid flow, such as density and flow velocity, can be recovered from the populations at any lattice point in the absence of *S_i_*:

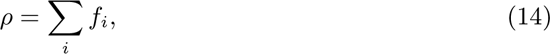

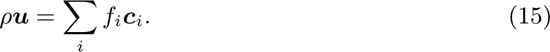

As detailed in [226], pressure and viscous stress can also be locally obtained from the populations. The link between the LB equation and the Navier-Stokes equation has been established through the Chapman-Enskog analysis [88, 227, 228].

Depending on the number of spatial dimensions and the number of discretised velocities, various lattice discretisations exist. For 2D problems, the most common stencil is D2Q9. In 3D, a wider range of stencils is available, most notably D3Q15, D3Q19 and D3Q27. A detailed discussion of velocity sets is given in [88]. Lattices with more velocities than nine in 2D and 27 in 3D have not been employed in IPMF applications. Although D3Q15 requires less memory and computational effort, it usually lacks accuracy and stability when compared with D3Q19 and D3Q27. For inertial flows, the D3Q27 lattice has shown its benefit in accuracy over 3D lattices with fewer speeds [229]. D3Q19 (Fig. 7) is the most common stencil employed for IPMF problems due to its compromise between numerical efficiency, accuracy and stability. We will comment on parameter selection strategies for IPMF simulations in Section 7.4.

**Figure 7:**
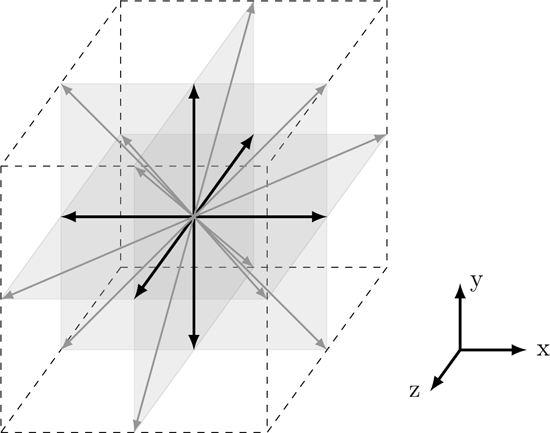
Illustration of the D3Q19 velocity stencil. The central lattice node is connected to 18 of its neighbours (indicated by the arrows) which are located on the principal planes (grey). Black arrows indicate velocity vectors along the main axes, grey arrows have two non-zero components. The enclosing cube (dashed) has an edge length of 2Δ*x*.

### 4.2 Equilibrium distribution and collision operators

A key step that led to the conceptual simplification of the LB equation was the replacement of the complex collision operator by much simpler relaxation-based operators [94]. In virtually all LB flavours that are currently used, the populations *f_i_* are relaxed to a local equilibrium state *f* ^eq^ (Section 4.2.1). This way, the collision operator in (Eq. 12) assumes the form

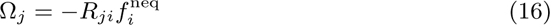

where **R** is a *q × q*-matrix that describes the relaxation process and *f* ^neq^ = *f_i_ − f* ^eq^, the non-equilibrium distribution, is the deviation of *f_i_* from its equilibrium *f* ^eq^. In IPMF applications, by far the most commonly used collision operator is the BhatnagarGross-Krook (BGK), also called single-relaxation time (SRT) collision operator, see Section 4.2.2. We comment on other relaxation operators in Section 4.2.3.

#### 4.2.1 Equilibrium distribution

The discretised equilibrium distribution *f* ^eq^ can be obtained from the Maxwell-Boltzmann distribution using either a truncated Hermite polynomial expansion or an expansion in Mach number [230]. The most commonly used equilibrium distribution is

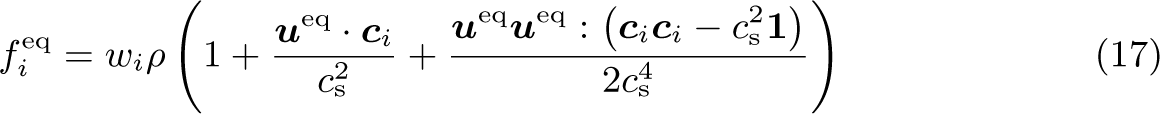

where **1** is the unit matrix. The quantities *w_i_* are the lattice weights, and *c*_s_ is the lattice speed of sound, both associated with the chosen lattice discretisation; see [88, 230, 231] for a detailed list of relevant parameters. The equilibrium velocity is given by

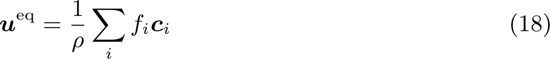

in the absence of forces (see Section 4.3 for changes due to the inclusion of external forces). The term in (Eq. 17) containing the quadratic expression *u*^eq^*u*^eq^ is necessary for the recovery of the advective term in the Navier-Stokes equation and therefore essential for all IPMF applications.

#### 4.2.2 BGK collision operator

In the widely used BGK or SRT model, the collision operator takes the form [225]

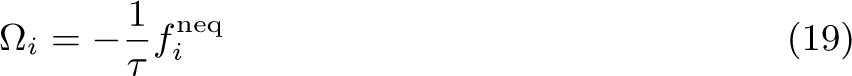

where *τ* is the single relaxation time. The dynamic viscosity of the fluid is linked to the BGK relaxation time as

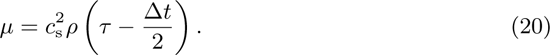

The BGK collision operator is extremely popular due to its simplicity and ease of implementation. However, the BGK model has limitations in terms of stability, errors arising from boundary conditions, and reaching very low or very large viscosity values [232, 233]. Although IPMF applications can usually be simulated with the BGK operator, some problems at higher Reynolds numbers can be overcome by choosing more sophisticated collision operators, such as MRT (see Section 4.2.3).

#### 4.2.3 Other collision operators

A few IPMF works have employed the multi-relaxation time (MRT) collision operator, rather than BGK [122, 135, 137, 176, 177]. The underlying idea behind the MRT collision operator is to use independent relaxation times (or frequencies) for different moments of the populations in order to improve the stability and accuracy of the method [232]. The *q* populations *f_i_* are mapped onto a *q*-dimensional moment space, and different moments *m_i_* (rather than populations *f_i_*) are relaxed with different frequencies *ω_j_* = 1*/τ_j_*. After relaxation, the moments are transformed back to the original population space. MRT enables the decoupling of bulk and shear viscosity parameters. This additional freedom is useful for low-viscosity problems and is therefore an advantage for IPMF applications. However, most IPMF problems can be safely simulated with the conceptually simpler BGK operator.

In recent years, various advanced collision operators have been proposed, such as the regularised [234, 235], entropic [236], cascaded [237] and cumulant [238] collision operators. Each of these operators comes with a set of advantages, mostly in terms of numerical accuracy and stability. In particular the entropic and cumulant collision operators are suitable for 3D turbulence modelling, but this advantage is usually not strongly visible at moderate inertia as found in IPMF applications. Since most of these advanced collision operators are more challenging to implement or more computationally expensive, they have not been employed for IPMF simulations.

### 4.3 Including external forces

External forces, including those from the immersed boundary method, usually enter the LB algorithm through the source term *S_i_* in (Eq. 12) and a modification of the equilibrium velocity ***u***^eq^ in (Eq. 18). There are various forcing schemes available since the forms of *S_i_* and ***u***^eq^ are not unique for a given physical force ***b***. This review does not aim to revise all existing forcing schemes. Instead, we present two popular methods, the Guo [239] and the Shan-Chen [240] forcing schemes, and refer to [241] for other schemes.

For the Guo forcing scheme [239], the source term takes the form

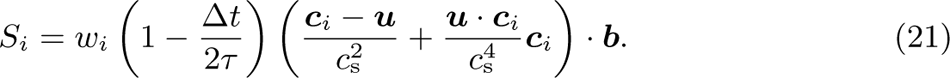

The equilibrium velocity is changed to

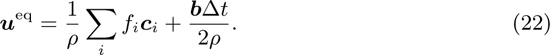

The Shan-Chen forcing scheme [240] is not to be confused with the Shan-Chen force that is often used to model multi-phase or multi-component flows. In the Shan-Chen forcing scheme, we have *S_i_* = 0 and

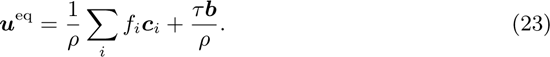

The Guo and Shan-Chen forcing schemes are second-order accurate in space and time under diffusive scaling when the macroscopic fluid velocity is additionally redefined as

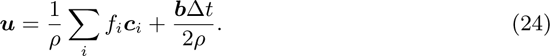

The Chapman-Enskog analysis shows that both schemes are equivalent in terms up to (*u*^2^) (see [88] for a detailed discussion).

There is a whole range of other forcing schemes that are also equivalent up to (*u*^2^) [241]. For IPMF applications, any second-order accurate forcing scheme is usually appropriate.

### 4.4 Non-Newtonian fluids

There is a growing interest in using non-Newtonian liquids in IPMF applications since non-Newtonian rheology gives rise to additional particle lift forces that interact with the inertial lift forces. The LB method in its original formulation recovers Newtonian fluid mechanics, but it can also be used for viscous non-Newtonian liquids and for viscoelastic liquids.

Several LB-based works have been published that considered inertial effects in combination with either power-law liquids [114, 136, 144, 145, 147, 148] or viscoelastic liquids [107, 149]. For viscous non-Newtonian liquids (such as shear-thinning or shear-thickening liquids), the strategy is to adapt the local relaxation time *τ* to the local strain rate to achieve the desired viscosity *via* (Eq. 20) [242, 243]. Viscoelastic liquids are more challenging to model since additional constitutive equations have to be solved. Details can be found, for example, in [244, 245].

## 5 Numerical methods for particles

Here we focus only on resolved particles with radii significantly larger than the fluid grid resolution, *a* Δ*x*, since under-resolved particle models in IPMF are still immature and need to be informed by resolved models. In order to simulate resolved particles in IPMF (see Section 3.2 for the physical particle models), their surface needs to be discretised, for example through mesh generation algorithms (Section 5.1). The numerical treatment of the particles is different for rigid and deformable particles (Section 5.2 and Section 5.3, respectively). The numerical coupling between particles and flow requires a separate discussion (Section 6.2).

### 5.1 Particle mesh discretisation

It is necessary to define and discretise the shape of the particles in order to resolve them in IPMF simulations. The most common approach for capturing the surface of a particle in IPMF simulations is to distribute multiple marker points over the surface of each particle. Depending on the type of the particle (rigid or soft) and the actual FSI algorithm chosen, the vertices need to be connected to their neighbours in order to create an unstructured surface mesh (Fig. 8). In most cases, such a mesh consists of *N_f_* flat triangular elements (or facets). Any pair of connected vertices defines the edge of two neighbouring triangles. For soft particles, the mesh is needed to numerically evaluate the local particle deformation and surface forces (Section 5.3). Table 2 summarises the cases for which a mesh is required.

**Figure 8:**
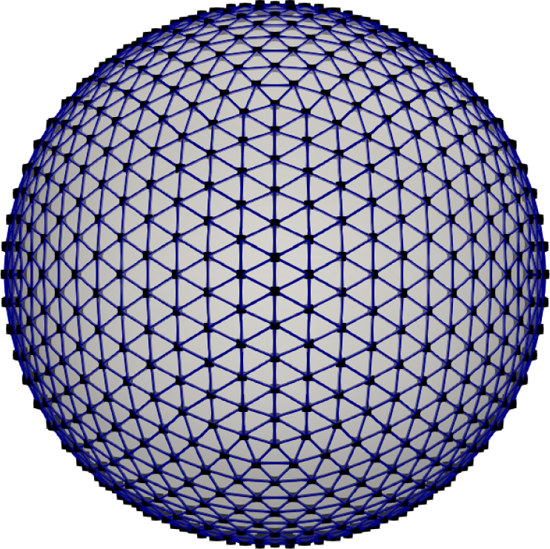
Surface markers (or vertices, black) and corresponding unstructured surface mesh (blue). The mesh consists of *N_f_* = 2420 elements and *N_v_* = 1212 vertices and was generated through the procedure presented in [246].

**Table 2:**
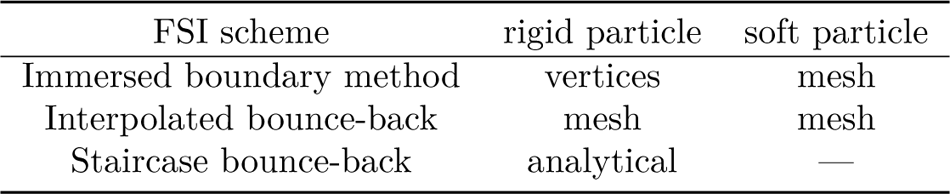
Discretisation requirements for rigid and soft particles and different FSI schemes. ‘vertices’: only surface points are needed; ‘mesh’: vertices with mesh are needed; ‘analytical’: analytically known particle shape is used, and no vertices are needed; ‘—’: not available or not practical. See Section 5.1 for the particle mesh discretisation and Section 6.2 for the numerical FSI schemes.

Distributing vertices and, if needed, generating a mesh is relatively straightforward for simple particle shapes, such as spheres, ellipsoids or red blood cells. Details of the surface mesh generation for spheres and red blood cells are provided in [246]. Although the vast majority of IPMF simulations involve simple particle shapes, open-source or commercial meshing software, such as CGAL [247] or Gmsh [248], might be used for more complex particle shapes.

A key consideration is the relative resolution of the particle discretisation compared to the lattice spacing, which can be quantified by the ratio *ℓ*^-^*/*Δ*x* where *ℓ*^-^ is the average distance between neighbouring vertices. As a rule of thumb, the numerical resolution of the particle mesh and the fluid lattice should be similar, *ℓ*^-^*/*Δ*x* 1, in particular, if the particle is soft and its deformation needs to be captured accurately. This rule is particularly important for most immersed boundary methods, although different algorithms might work best with different ratios. If the interpolated bounce-back method is used, *ℓ*^-^*/*Δ*x* is less constrained, and the particle discretisation is largely determined by the degree of complexity of the particle shape and the particle deformation expected during the simulation. Section 6.2 provides more details about the different FSI schemes employed in IPMF.

In some cases, it is also useful or necessary to create a particle volume mesh [249] or a different internal particle structure [250], although these approaches have not yet been used for IPMF problems. Interior vertices are required by some immersed boundary methods or when the particle has an internal structure that determines the deformation of the particle, see Section 6.2.2.

### 5.2 Rigid particles

Rigid particles are characterised by the constant distance between any pair of marker points on their surface. Therefore, the rigid particle algorithm should translate and rotate the particles according to (Eq. 5) and (6) while keeping the particles’ shape invariant. An example algorithm for the simulation of rigid particles in inertial microfluidics can be found in [107].

Several numerical schemes can be used for the implementation of the motion of rigid particles. However, when choosing an algorithm, its numerical stability relative to that of the fluid solver is critical. Ideally, the rigid-particle solver should have similar stability to the fluid solver over an extensive range of time step sizes. A good candidate is the Verlet integrator [251, 252]: it provides second-order accuracy and good numerical stability. Additionally, it preserves the time reversibility and the symplectic form of the governing equations. Symplectic solvers are a class of numerical algorithms which, by construction, ensure that the system’s total energy is conserved during numerical integration [253].

The representation of the orientation of the particles in 3D requires special attention. Due to the properties of the Lie group SO(3), which represents the space of all possible 3D rotations, it is not possible to describe the orientation of a particle without the emergence of singularities using 3D vectors [254]. These singularities are commonly called ‘gimbal lock’. The gimbal lock can be avoided by using either a rotation matrix or unit quaternions to represent the orientation of the particles. Quaternions are a 4D extension of complex numbers, and their mathematical multiplication rules can encode SO(3) without encountering singularities [255]. Quaternions have several advantages over rotation matrices in numerical schemes for rigid particles; they have a smaller memory footprint than matrices, and they are more algorithmically efficient when performing rotation operations on vectors. Detailed information on the implementation of quaternions can be found in [256].

### 5.3 Soft particles

To treat soft particles numerically, we need to discretise the physical model presented in Section 3.2.3. We distinguish between the outer membrane of soft particles (Section 5.3.1) and their internal properties (Section 5.3.2).

#### 5.3.1 Hyperelastic model for membrane-based particles

For hyperelastic particles that are made of a thin membrane (*e.g.*, RBCs, capsules, vesicles), numerical treatments of in-plane elasticity, bending, and — if required — surface and volume conservation are needed. In the following we will discuss a commonly used approach based on the finite-element method in more detail [246]. We will not cover the lattice spring method (LSM) here since it has been used only in a small number of IPMF works [70, 102, 122, 147]. Viscoelastic membrane models are covered, for example, in [209, 210], although they have not yet been used for IPMF applications.

The elastic and bending energies of a soft particle, (Eq. 8) and (Eq. 9), may be discretised as [191]

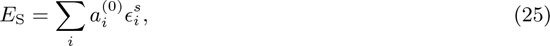

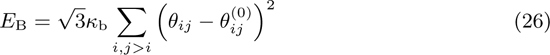

where the sum in (Eq. 25) runs over all surface elements *i* and the sum in (Eq. 26) runs over each pair of neighbouring elements, *a*^(0)^ is the area of the undeformed element, *E^s^* is the strain energy density of the deformed element, *θ_ij_* is the current angle between the normal vectors of two neighbouring elements, and *θ*^(0)^ is the angle between the normal vectors of two neighbouring elements of the undeformed mesh. The strain energy density of each element, *E^s^*, can be calculated through a finite-element-based scheme using linear shape functions [246]. See [257] for a detailed discussion of alternative bending models. In cases where the enclosed volume or the surface area of the membrane is constant, the constraints on volume and surface area can be implemented using penalty energy terms [191]:

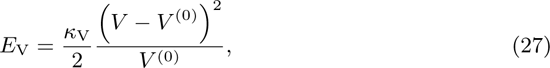

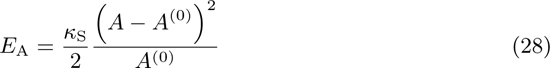

where *κ*_V_ and *κ*_S_ are the corresponding penalty moduli, *V* and *A* are the current volume and surface area, and *V* ^(0)^ and *A*^(0)^ are the volume and surface area of the undeformed particle.

The force acting on each vertex *i* of the particle surface can be obtained from the total particle energy *E* = *E*_S_ + *E*_B_ + *E*_V_ + *E*_A_ using the principle of virtual work [213, 214]

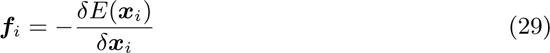

where ***f****_i_* can be written directly as a function of vertex positions, ***x****_i_*, see [258] for implementation details. Fig. 9 demonstrates the capabilities of the model.

**Figure 9:**
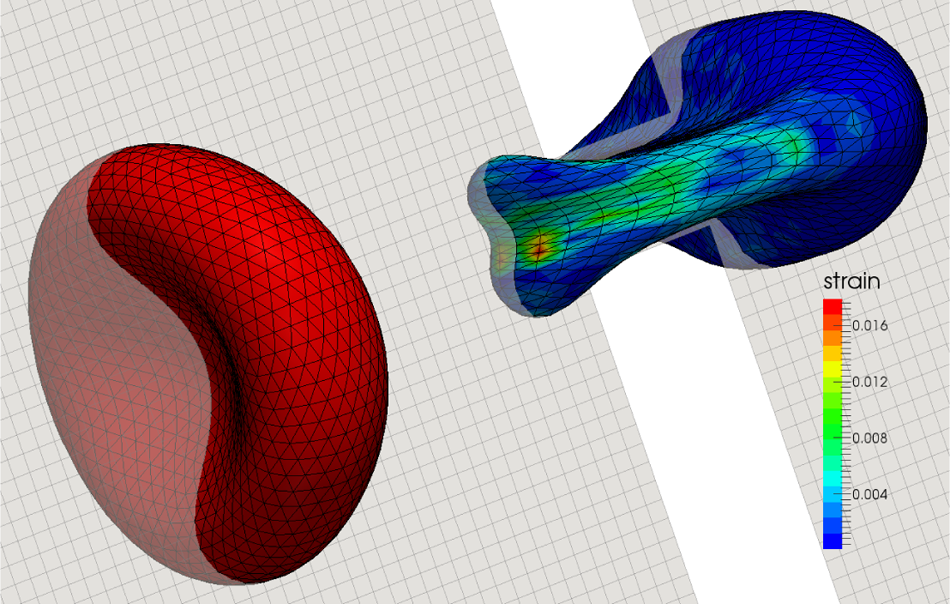
Visualisation of a single RBC squeezing through a spleen slit. The mesh of the undeformed RBC (left) and the deformed RBC (right) is indicated by black lines. One plane of the fluid grid is shown in grey. The colour of the deformed RBC indicates the strain force (N) arising from local relative surface deformation.

#### 5.3.2 Internal particle properties

Some particles have an internal structure or are filled with a liquid whose properties are different from those of the suspending liquid. For these types of particles, additional numerical methods are needed. At the time of writing this review article, no LB-based IPMF paper has been published in which soft particles with complex internal structures have been considered. However, we anticipate that the behaviour of nucleated cells or similar particles in IPMF devices will be simulated in the near future.

The interior viscosity of red blood cells, vesicles and capsules is often different from that of the suspending liquid. In practice, the fluid nodes inside the membrane need to be tracked and updated as the particle deforms and moves. Different strategies have been suggested, such as simple ray-casting [259] and the Hoshen-Kopelman algorithm [260]. More recently, a fast tracking algorithm has been proposed by computing the scalar product of area-weighted surface normals and local distance vectors in the vicinity of the membrane [261]. Once each lattice node knows its viscosity *µ*(***x***), the local lattice-Boltzmann relaxation time is calculated *via* (Eq. 20).

For eukaryotes (*i.e.*, biological cells with a nucleus), a compound membrane model may be needed where the outer cell membrane and the inner nuclear envelope are interconnected by a cytoskeleton within the cytoplasm. The nuclear envelope can be modelled and discretised in a way similar to the RBC membrane model, and the cytoskeleton can be represented as cross-linked filaments consisting of discrete particles [250] or cylindrical segments [262].

## 6 Numerical boundary conditions and fluid-structure interaction

In this section, we present the boundary conditions and FSI schemes commonly used for LB-based IPMF applications. In Section 6.1, we revisit the different types of boundary conditions identified in Section 3.3 from a numerical point of view. Since FSI is typically the biggest numerical challenge in IPMF, we cover it in more detail in Section 6.2.

### 6.1 Boundary conditions

In Section 3.3, we distinguished between three different types of boundary conditions: 1) device-fluid, 2) particle-fluid, and 3) inlet and outlet boundary conditions (see Fig. 6). Since the first and third types do not require FSIs and are relatively straightforward to implement, we outline them in Section 6.1.1 and Section 6.1.2, respectively.

#### 6.1.1 Device-fluid boundaries

Numerical algorithms for the device-fluid boundaries are well established and do not pose significant challenges for IPMF simulations. The easiest method for realising the no-slip condition at stationary walls in LB-based simulations is the simple (or halfway) bounce-back (SBB) method (Fig. 10). In SBB, a post-collision population *f\**streaming from a fluid node ***x***_f_ to a solid node ***x***_s_ is simply reflected (bounced back) to its original node ***x***_f_ while reversing its direction [100]:

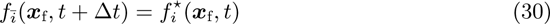

**Figure 10:**
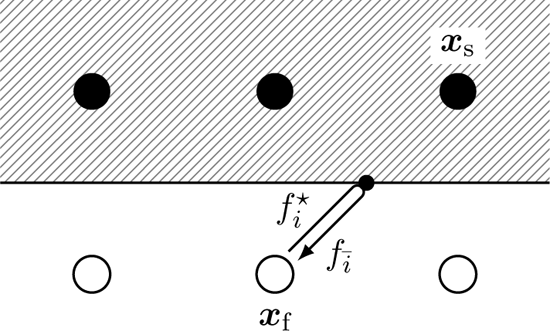
Illustration of the simple bounce-back (SBB) algorithm. A post-collision population *f ^*^* that would stream from a fluid node ***x***_f_ to a solid node ***x***_s_ bounces back when it reaches the wall (solid line) which is located half-way between the fluid and solid nodes. The point where the bounce-back event occurs is indicated by a small black dot. The population returns to its starting point as *ℓ*^-^.

where ^-^*i* is defined through ***c***_-*i*_ = ***c****_i_*. In this scheme, the no-slip condition is recovered to second-order accuracy if the physical wall is flat, aligned with one of the principal lattice axes, and located halfway between ***x***_f_ and ***x***_s_. SBB is a method that is only available for LB or similar methods since the populations do not exist in macroscopic CFD approaches, such as finite volume or finite difference methods. A clear advantage of SBB is its local character and, thus, relative ease of parallelisation. In situations where the geometry is not aligned with the lattice axes, SBB leads to a ‘staircase’ representation of the boundary, and the method’s accuracy degrades to first order [263].

A common improvement of the SBB for curved boundaries is the interpolated bounce-back (IBB) method, where the distance between the lattice nodes and the actual wall location is taken into account. The Bouzidi method [264] is a popular IBB variant that uses two or three neighbour nodes to obtain a second-order representation of curved boundaries *via* linear or quadratic interpolation at the population level. IBB methods are often less local than SBB, potentially creating challenges for parallelisation and for moving particles (see Section 6.2.1). For example, the Bouzidi method requires at least two fluid lattice nodes between nearby boundaries. This limitation of the Bouzidi method can be overcome by approximating the equilibrium and non-equilibrium parts of *f_i_* at a fictitious node at the exact boundary location, and carrying out interpolations with only the immediate boundary lattice node [265, 266].

Bounce-back methods can be extended to situations where the boundary is moving, either as an imposed movement (*e.g.*, the lid in a Couette flow) or as part of fully coupled fluid-structured interaction (Section 6.2.1). The resulting momentum exchange at a moving wall is captured by a correction term, here shown for SBB [100]:

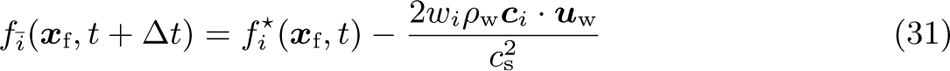

where ***u***_w_ and *ρ*_w_ are the velocity of the wall and the density at the point of intersection between the lattice link and the wall, respectively. For IBB, the correction term has to be applied to the fraction of the post-collision population that streams into the wall boundary [267].

All bounce-back-based boundary conditions show a dependency of the exact wall location on the chosen value of the relaxation time *τ* when the BGK collision operator is used. This problem usually worsens for large values of *τ* and is, therefore, not a common problem in IPMF applications where the viscosity (and therefore *τ*) is relatively small. Replacing the BGK by the MRT or other collision operators with well-chosen relaxation times can avoid this unphysical dependency [268]. A detailed analysis of the bounce-back method and strategies for improving its accuracy are given in [88].

Bounce-back methods are classified as link-wise boundary conditions since the wall location is somewhere between the fluid and solid nodes. There exists a large range of alternative LB boundary conditions where the boundary condition is enforced directly on lattice nodes, *e.g.* [269–271]. These methods are often called ‘wet-node’ approaches. Since wet-node methods are rarely used in IPMF applications, we do not cover them here.

Finally, it is also possible to use the immersed-boundary (IB) method for the surface of the geometry. However, since the IB method is more commonly employed for FSI problems, we cover it in Section 6.2.2 instead.

#### 6.1.2 Inlet and outlet boundaries

Nearly all published papers with LB-based IPMF simulations use periodic boundary conditions to treat the channel inlet and outlet. Periodic boundary conditions are straight-forward to implement, both for the fluid and for the particles. In particular, particles do not have to be created or removed once the simulation has started, see Section 7.2.

To drive the flow in a geometry with streamwise periodic boundary conditions, there are two strategies: 1) use a body force ***b*** or 2) superimpose a pressure drop Δ*p* on the periodic boundary condition. The first strategy is particularly suitable for straight channels where the pressure gradient is essentially constant and can be replaced by a constant body force ***b*** = *p*. The pressure fluctuations caused by the presence of the particles are then automatically captured by the LB algorithm. For more complex geometries, it is often more suitable to impose an overall pressure drop Δ*p* between the inlet and outlet planes [272]. The LB algorithm will then recover the correct pressure field in the interior of the domain, including any pressure fluctuations caused by the particles.

Although tempting and conceptually simple, periodic boundaries do not come without their own problems. Since an infinite array of particles is simulated, long-range particle-particle interactions across periodic boundaries need to be controlled by having sufficiently long channel segments subject to sensitivity tests of the chosen unit cell length [117, 273].

There are situations where periodic boundary conditions are not suitable, for example, if the geometry of interest cannot be approximated by a periodic unit cell. One example is the cross-slot junction with two inlets and two outlets simulated in [131]. In these situations, it is common to impose a fully developed velocity profile at the inlet and a pressure condition at the outlet, or the opposite. The most frequently used LB scheme in such cases is the non-equilibrium extrapolation method [274].

Challenges remain in developing more suitable boundary conditions for complex geometries. In particular, in inertial flows, flow field distortions can propagate far down-stream. By imposing a fully developed velocity profile at the inlet, any upstream influence is neglected, which might make the simulation unsuitable. Pressure waves and flow distortions leaving the domain should not be reflected back at the outlet, which imposes additional constraints on the performance of the chosen numerical scheme. Concluding, more work is needed to enable accurate and practical LB simulations of IPMF device segments that cannot be approximated as periodically repeating units.

### 6.2 Fluid-structure interaction methods

The purpose of the FSI algorithm is to impose appropriate boundary conditions on the moving particles and to evaluate forces and torques that accelerate the particles in flow. In LB-based IPMF applications, there are different algorithms used over the years, some of which are suitable for either rigid or soft particles, or both. The key point is that any re-meshing of the fluid domain is undesired as the original LB method relies on a fixed lattice. Therefore, all FSI schemes covered here are based on a fixed fluid lattice and moving particle meshes which need to be coupled. Here we will focus on the two most commonly employed methods in LB-based IPMF: 1) bounce-back-based methods that operate on the level of the populations and involve a momentum-exchange algorithm (MEA) (Section 6.2.1) and 2) the immersed boundary (IB) method that uses forces to mimic the existence of boundaries (Section 6.2.2). Other FSI methods are less established in IPMF, such as the external boundary force method [275] and the Noble-Torczynski method [173]; we do not cover them here.

#### 6.2.1 Bounce-back methods and momentum exchange algorithm

The bounce-back-based algorithms for FSI are essentially the same as those in Section 6.1.1. In order to calculate the force and torque the suspended particles experience, a MEA is required. Additionally, the location of the boundary changes every time step, which requires the fluid-solid links to be updated dynamically and some nodes to be switched from fluid to solid, or vice versa.

In the vast majority of cases, SBB and IBB are used for rigid particles. It is possible to employ bounce-back-based methods for soft particles, although instead the IB method (Section 6.2.2) is normally the method of choice in those cases. We found only two studies where IBB has been used for soft particles in IPMF [102, 147].

The MEA treats populations crossing fluid-particle boundaries as discrete mass packets that exchange momentum with the particle surface [276]. The momentum that is transferred from the fluid to the solid due to a single population initially moving in direction ***c****_i_* and then bouncing back at point ***x***_b_*_i_* on the boundary is (in 3D)

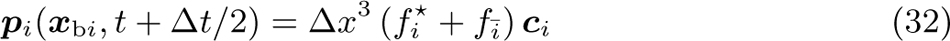

where *f^*^_i_* is the post-collision population moving towards the boundary and *ℓ*_-_ is the population moving in the opposite direction after the bounce-back event. For SBB, *ℓ*_-*i*_ is given by (Eq. 31). Since the population bounces back between time steps, the force is evaluated at half-time steps, *t* + Δ*t/*2. (Eq. 32) holds for simple and interpolated bounce-back methods. The total force and torque acting on a particle at *t* + Δ*t/*2 are calculated as

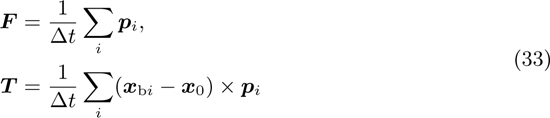

where the sum runs over all links on which a bounce-back event occurs and ***x***_0_ is the centre of mass of the particle.

Two fundamentally different strategies have been suggested to treat the interior of rigid particles and the fresh nodes that are crossed by the moving particle surface. Ladd [100] kept the fluid inside the particles and included the internal stresses by applying the MEA both to the exterior and interior fluid regions. This approach will be problematic if inertia is significant since the inertia of the internal fluid can affect the dynamics of the particle. However, the fresh node treatment consists of merely passing nodes through the surface without further modification. Aidun and Lu [277] proposed a different approach where the MEA only considers the exterior fluid. Different fresh-node strategies have been proposed [278, 279].

Table 2 reminds us that SBB does not usually require meshing of the particle surface since SBB is mostly applied when particles have simple shapes (*e.g.*, spheres or ellipsoids) and the identity of fluid and solid nodes can be easily established at each time step. IBB, however, requires detailed information about the location of the bounce-back event along the link between lattice nodes, in particular when particles are deformable. A particle surface mesh combined with a ray-tracing algorithm [280] can be employed for this purpose. Particular attention must be given to the relative resolutions of the fluid lattice and the particle surface mesh [281].

Bounce-back-based approaches do not generally conserve mass when particles are moving. However, the MEA emerges naturally from the bounce-back scheme, and no additional computation is needed to obtain traction vectors from the surrounding flow field. The MEA employing (Eq. 32) has been shown to violate Galilean invariance on order (*u*^2^), and a correction term has been proposed to remove this artefact [282]. While the SBB is simpler to implement than IBB, it suffers from a staircase approximation of the moving particles. IBB has second-order accuracy but is less local and more cumbersome to implement.

In conclusion, the implementation of bounce-back-based FSI schemes for IPMF requires several careful considerations, but the available methods are well established and of sufficient efficiency and accuracy.

#### 6.2.2 Immersed boundary method

The second class of FSI methods commonly used for IPMF is the IB method in its various flavours. Peskin’s underlying idea was to use body forces to manipulate the fluid flow field around the boundary to satisfy the no-slip condition at the boundary [283, 284]. There are versions of the IB method coupled to an LB solver for both soft [246, 285–288] and rigid particles [289–294]. It is not possible to cover all flavours of the IB method in detail here. Instead, we provide a concise outline and refer to recent review articles [295, 296].

The general IB algorithm consists of a few key steps (not necessarily in this order):

- Interpolate the fluid velocity ***u***(***X***) at the position ***x****_i_* of each particle mesh vertex *i* (see Section 5.1 for details about the mesh).
- Calculate forces ***f****_i_* acting on each vertex *i*. See Section 5.3 for the force calculation in the case of soft particles. See below for rigid particles.
- Spread vertex forces ***f****_i_* to the fluid where they will be treated as body forces ***b***(***X***) according to Section 4.3.
- For each particle in the simulation, update its position and orientation. See Section 5.2 for rigid particles. See below for soft particles.

Here we distinguish between position vectors ***X*** of fluid nodes on the regular Eulerian lattice and position vectors ***x****_i_* denoting the location of a Lagrangian vertex with index *i*.

Fig. 11 illustrates the interpolation and spreading steps. In *d* spatial dimensions, the discretised forms of the velocity interpolation and the force spreading steps, which are key to any IB algorithm, can be written as [284]

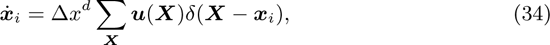

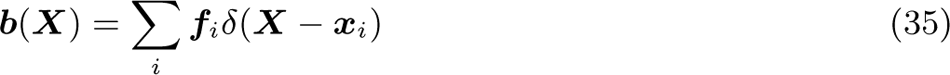

where ***x*** *_i_* is the interpolated velocity of vertex *i* and *δ*(***X x****_i_*) is a discrete delta distribution with the dimension of *L^−d^* where *L* is length. Note that ***ℓ****_i_* has the dimension of a force and ***b*** has the dimension of a force per *L^d^*.

**Figure 11:**
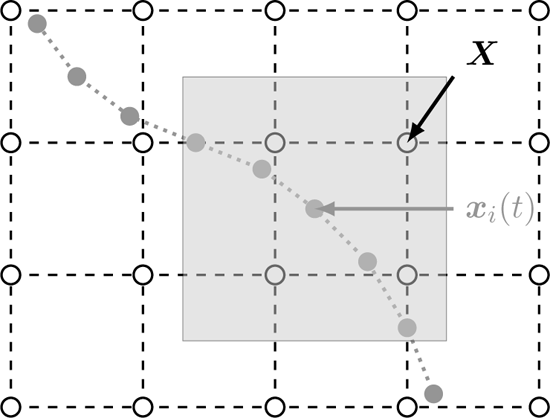
Illustration of interpolation and spreading in the immersed boundary method. The velocity of the Eulerian lattice nodes (open circles, coordinates ***X***) is interpolated at the location ***x****_i_*(*t*) of each Lagrangian vertex (solid circles), see (Eq. 34). Only lattice nodes within the interpolation window (grey square) around the Lagrangian vertex of interest are considered. Force spreading, (Eq. 35), works the other way around, where each Lagrangian vertex distributes its force to the lattice nodes within the spreading window.

A key feature of the IB method is the shape of the discrete delta distribution *δ*(***X − x****_i_*). An important simplification is the factorisation *δ*(***x***) = *φ*(*x^t^*)*φ*(*y^t^*)*φ*(*z^t^*)*/*Δ*x*^3^ in 3D, or *δ*(***x***) = *φ*(*x^t^*)*φ*(*y^t^*)*/*Δ*x*^2^ in 2D, where *φ*(*x^t^*) is a suitable 1D kernel function and ***x****^t^* = ***x****/*Δ*x* = (*x, y, z*)*^T^ /*Δ*x*. The most commonly used forms are

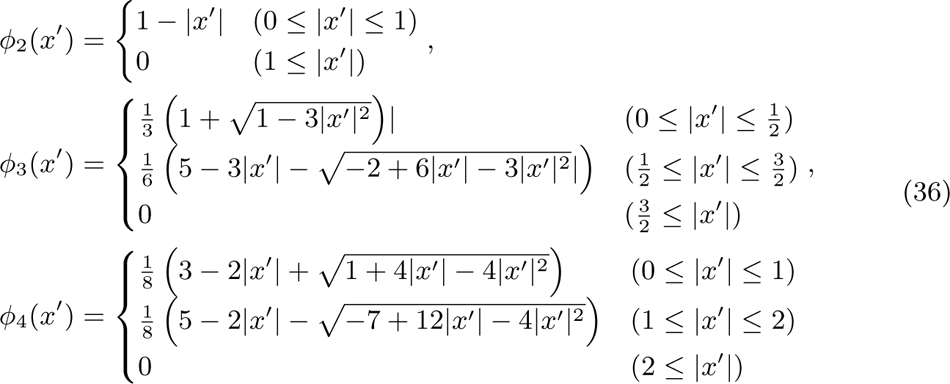

where *φ*_2_, *φ*_3_ and *φ*_4_ are often called the 2-point, 3-point and 4-point stencils since they cover two, three or four grid points along each coordinate axis, respectively (Fig. 12). Therefore, IB interpolation and spreading for each Lagrangian vertex involves a square or cube covering *n^d^* lattice nodes where *n* is the width of the stencil (*n* = 2, 3, 4). The rationale behind these forms and mathematical derivations are available elsewhere [284]. There is an important difference between the IB algorithms for soft and rigid particles.

**Figure 12:**
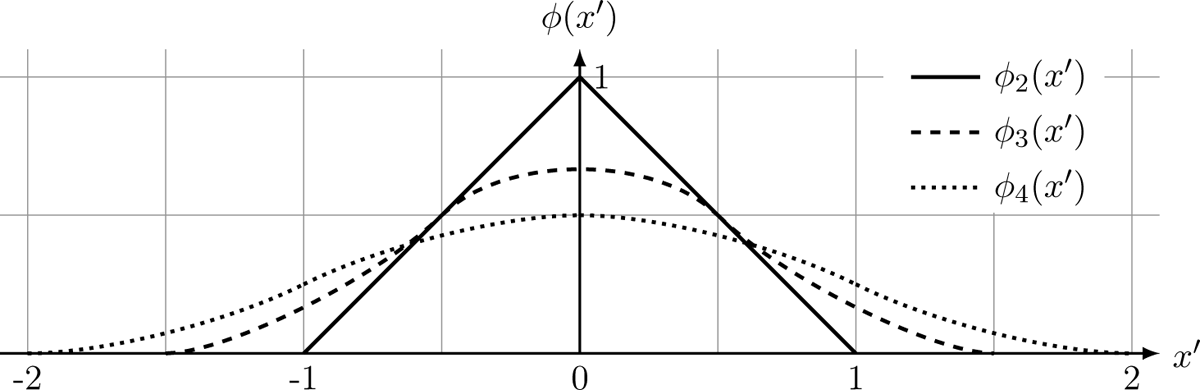
Plots of the interpolation stencils in (Eq. 36). The stencil *φ_n_* covers *n* lattice nodes along each coordinate axis.

For soft particles, the vertex velocity is determined by (Eq. 34) and particle deformation is caused by different vertices, *i* and *j*, generally moving in a way that the distance ***x****_i_* ***x****_j_* changes with time. The resulting deformation of the mesh elements then leads to forces acting on each vertex, for example, *via* (Eq. 29). These forces are then spread to the Eulerian lattice through (Eq. 35) after which they enter the LB algorithm (Section 4.3). Note that, in general, the vertices used for the force calculation and the IB interpolation and spreading do not have to be the same [297, 298], although in most works both are identical. Vertex positions are usually updated using a forward-Euler approach:

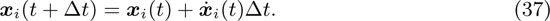

The IB algorithm for soft particles is detailed in [107, 246, 299, 300].

There exist various different IB algorithms for rigid particles. The challenge in applying the IB method to rigid particles lies in satisfying the rigidity conditions ***x****_i_* ***x****_j_* = const and the no-slip conditions in (Eq. 34) simultaneously. The vertex forces ***f****_i_* need to be calculated in such a way that these conditions are met to a sufficient level of accuracy. Several IB flavours have been used for rigid particles in combination with the LB method:

- Feng and Michaelides’s direct-forcing method [290] applies an explicit correction due to the mismatch of desired and interpolated velocities at each Lagrangian vertex. Although relatively simple to implement, this method does not strictly satisfy the no-slip condition at the particle surface.
- The implicit IB method [291] solves the system of equations resulting from the simultaneous satisfaction of no-slip and rigidity conditions. Since this method requires inverting large matrices at every time step, only few works have adopted this approach.
- Multi-direct-forcing methods perform multiple iterations during each time step to approximate the solution of the implicit method [292–294]. This approach combines the ease of implementation of the direct-forcing method and the accuracy of the implicit method.

For details of the algorithms and implementation strategies, we refer the reader to the original publications.

An important consideration for any IB method is the average distance between neighbouring mesh vertices, *ℓ*^-^, in relation to the lattice spacing, Δ*x* (see Section 5.1). If *ℓ*^-^*/*Δ*x* is significantly larger than unity, there are ‘holes’ in the mesh and fluid can leak through the mesh surface. This detrimental effect starts to become visible for *ℓ*^-^*/*Δ*x* 2 [246]. At the other end, if *ℓ*^-^*/*Δ*x* is too small, neighbouring mesh vertices see approximately the same flow field since the velocities ***x*** *_i_* are obtained through interpolations in (Eq. 34). Thus, it is often recommended to use a relative spacing of *ℓ*^-^*/*Δ*x* 1 [246]. As a consequence, different meshes need to be prepared for the same particle at different spatial resolutions.

## 7 Additional considerations

IPMF simulations come with a range of additional requirements that have not been covered in the previous sections. Here we will address particle interaction forces (Section 7.1), strategies to initialise simulations (Section 7.2), code parallelisation and grid refinement (Section 7.3), and strategies for parameter selection (Section 7.4).

### 7.1 Particle-particle interaction forces

As long as particle surfaces are a few lattice points away from each other, the particle interactions are hydrodynamic and can be accurately handled by the LB and FSI algorithms without additional considerations. However, when particles come close (*e.g.*, in denser suspensions), the lattice resolution may not be sufficient to capture lubrication forces and avoid particle overlap.

If particles are rigid and circular or spherical, conventional lubrication forces or elastic repulsion forces can be applied based on the centre-to-centre distance between the particles. For example, Başağaoğlu *et al.* [130] used a Lennard-Jones potential. Liu and Wu [172, 173] employed a discrete-element method where contact forces have both elastic and damping contributions. Schaaf *et al.* [106] followed a different strategy by implementing an event-based Euler step to handle particle collisions. The collision treatment in several other works [148, 159, 165, 169] is based on the short-range repulsion model proposed by Glowinski *et al.* [301]. Liu *et al.* [161] employed Wan and Turek’s repulsion force model [302].

For more complex particle shapes and deformable particles, forces are usually calculated for any pair of nearby surface vertices belonging to two different particles. Hu *et al.* [141] used a velocity-dependent lubrication force for rigid circular and ellipsoidal particles, following Ding and Aidun [303]. Similar approaches have been used for soft particles, although mostly in non-inertial settings [304, 305]. More work is required to accurately simulate the contact dynamics in denser suspensions of soft particles in IPMF.

### 7.2 Simulation initialisation, particle insertion, and removal

Due to the movement of particles, all IPMF applications are transient, and initial conditions can play an important role. However, the background flow field in the absence of particles is steady in most cases, which allows for relatively straightforward initialisation strategies.

The simplest approach to initialise the background flow is to start a simulation with zero velocity and wait until the flow field has converged under the imposed boundary conditions or driving force. If the simulation program supports a checkpoint functionality, the converged state of a simulation can be stored to the hard drive and used as the initial condition for other simulations. Initialisation strategies for LB simulations have been discussed previously, *e.g.* [269, 306–310].

Assuming that the background flow field has already been established, particles may be ‘dropped’ in the simulation domain subsequently, although the sudden addition of particles will lead to pressure waves and changes in the flow field. Since the flow requires some time to adjust to the presence of particles, the first few hundred or thousand time steps are usually unphysical and should be excluded from the data analysis.

It is not always obvious at which positions particles should be initialised. For example, when a subset of an IPMF device is simulated (see Section 3.3 and Section 6.1.2), it is generally unknown at which position and under which deformed state particles would enter the numerical domain. Assuming that the channel upstream of the simulated segment is straight and long, a good strategy is to obtain the lateral equilibrium positions of the particles in separate simulations of straight channels and then initialise particles at these positions when they enter the domain of interest. Two migration characteristics can be exploited to accelerate the straight channel simulations. First, lateral particle migration is faster for particles initially positioned between the equilibrium position and the channel wall than for particles initially positioned between the equilibrium position and the channel centre. Second, particle migration in the radial direction is significantly faster than in the circumferential direction [311].

The initialisation of particles in periodic domains is usually straightforward. Depending on the aim of the study, the desired number of particles can be initialised at predefined or random positions. In the latter case, particle-particle and particle-wall overlap checks may be necessary. For denser suspensions, it can be useful to initialise particles with a reduced size and run a pre-simulation to grow particles to their full size [312].

Most LB-based IPMF studies have employed periodic boundary conditions. We are not aware of LB-based IPMF studies where particles are continuously inserted at the inlet and removed at the outlet (see [305, 313, 314] for non-inertial applications of inflow boundary conditions for red blood cells). There is a need for more advanced particle initialisation strategies that faithfully capture the upstream behaviour of the system when subsets of non-periodic geometries are simulated and the assumption of a long channel is not appropriate.

### 7.3 Parallelisation and grid refinement strategies

Accurate IPMF simulations typically demand a large number of grid points due to the requirement of high resolution. Since inertial effects are long-ranged, it is normally necessary to simulate domains that are much larger than the particles inside. There are two fundamental strategies to address this issue: code parallelisation and local grid refinement.

While the LB algorithm can be easily parallelised, the parallelisation of the moving particles and the associated FSI treatment is more challenging. Parallel open-source LB codes are available, *e.g.* [222–224, 315–318]. Detailed parallelisation strategies for particle-laden LB simulations have been published previously [288, 312, 314, 319–322]. Recent breakthroughs in the parallelisation of dissipative particle dynamics codes [323–327] also provide insights for further improving the parallel performance of existing LB codes in simulating particle-laden flows. Particularly noteworthy is lbmpy [328], a metaprogramming system for automatic code generation for parallel LB simulations.

For IPMF problems, the flow field around the particles usually shows finer features than in the regions farther away from the particles. Therefore, it should be possible to have a more refined fluid region around the particles and a coarser mesh elsewhere. Local grid refinement for LB simulations is an active research field, *e.g.*, [329–334]. However, dynamic grid refinement around moving particles using the LB method is largely unexplored.

### 7.4 Parameter selection

In order to set up and run IPMF simulations that are accurate, stable, and efficient, there are a few helpful rules of thumb, ideally considered in this order:

1. Define the geometry for the simulation first. If periodic boundary conditions are used, undesired interactions of periodic images of particles need to be avoided. As a guideline, start with a periodic system length of around ten particle diameters and test whether results are independent when this length is varied by several particle diameters. The domain length should be sufficiently large to ensure that periodic images have no effect; the minimum domain length depends on particle-to-channel confinement, Reynolds number, and particle concentration.
2. Once the geometry is defined, choose a lattice resolution that balances accuracy and efficiency. The numerical algorithms chosen require a certain number of grid points to resolve the particle diameter (a good starting point is around 10Δ*x*). A challenge of IPMF modelling is that particles are often small compared to the channel size. Therefore, it is advisable to perform a grid-independence study and use the lowest resolution that still gives sufficiently accurate results. Example grid-independence studies can be found in [117, 273].
3. Based on the chosen spatial resolution, the desired Reynolds number dictates the range of fluid viscosity *ν* and characteristic velocity *U via* (Eq. 3). Keep in mind that *U* in units of Δ*x/*Δ*t* should ideally not exceed 0.1 and that the relaxation time *τ* in units of Δ*t* should not get too close to 0.5. As a starting point, use *τ* ≈ Δ*t*. Generally, reducing *τ* (and therefore *µ*), while keeping the spatial resolution and Reynolds number fixed, reduces the velocity *U* and therefore the time step Δ*t*. This variation leads to a trade-off between efficiency on the one hand (smaller Δ*t* means more time steps need to be simulated for the same physical time) and accuracy and stability on the other hand (smaller Δ*t* usually means more stable and accurate results, but *τ/*Δ*t* → 0.5 can cause numerical instability). An in-depth discussion of the choice of parameters in LB simulations can be found in [88].
4. In case the flow is driven by prescribed inlet and outlet conditions, rather than by periodic boundary conditions and a driving force, a pressure drop between the inlet and outlet is required. Since pressure in LB simulations is linked to the variation of the fluid density, it is important that the resulting pressure drop does not lead to unreasonable density variations along the flow axis. In order to balance the numerical pressure difference between the inlet (*p*_in_) and the outlet (*p*_out_) with other numerical requirements, it is worth keeping in mind the Hagen-Poiseuille law for the laminar flow (average velocity *u*-) in a straight tube with radius *R* and length *L*, which is a good starting point even for different channel shapes:

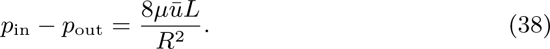

The resulting density difference is *ρ*_in_ − *ρ*_out_ = (*p*_in_ − *p*_out_)*/c*^2^ and, as a rule of thumb, should not exceed a few percent of the average density.

## 8 Example Cases

This section provides four example cases that capture the general physics of IPMF: the lateral migration of a rigid and a soft particle in a square duct (section 8.1 and section 8.2, respectively), the interaction of a pair of soft particles in simple shear flow (section 8.3), and the formation of a train of soft particles in a square duct (section 8.4). We use the in-house BioFM code and compare results with existing literature data. For all example cases, we use the D3Q19 lattice [225], the BGK collision operator [335] with relaxation time *τ* and the forcing method of Guo *et al.* [336]. Throughout this section, we report the kinematic viscosity *ν* = *µ/ρ*, rather than the dynamic viscosity of the liquid.

### 8.1 Example case 1: migration of a single rigid particle in a square duct

The first example case simulates the lateral migration and equilibrium position of a rigid, spherical, neutrally buoyant particle placed in a straight duct with square cross-section with width 2*w*. Originally proposed by Lashgari *et al.* [273], the trajectories and equilibrium positions are compared for three different confinement values, *χ* = *a/w*. The confinement is varied by modifying the particle radius *a*, resulting in *χ* = 0.1, 0.2 and 0.287.

Fig. 13(a) and (b) show a full 3D and a 2D cross-sectional schematic of the geometry. Table 3 contains values of the relevant fluid and particle properties. For this case, we follow Lashgari *et al.* [273] by defining the Reynolds number as Re = *U* 2*w/ν* where *U* is the mean cross-sectional flow velocity. The flow is driven by a body force to reach the desired velocity *U*. Simulations are initialised by dropping the particles in the simulation box and then driving the flow, starting at *t* = 0.

**Figure 13:**
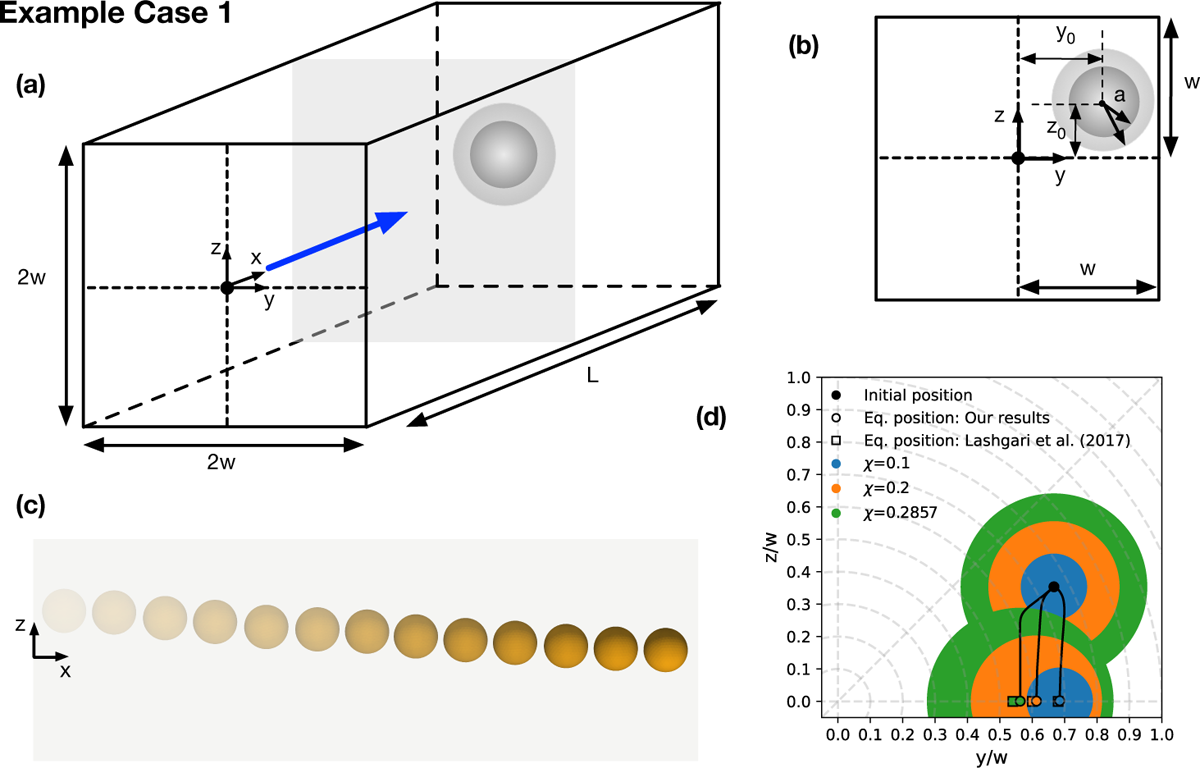
Example case 1: migration of a single rigid particle in a square duct. (a) 3D schematic. The grey plane denotes the channel cross-section. Particles of different sizes are illustrated by the two spheres. (b) 2D cross-sectional schematic. (b) Migration path of a particle with *χ* = 0.2 along the *z*-axis; multiple time instances are overlaid with higher saturation indicating later time. Note that the axial direction is not to scale. (d) Lateral migration paths for different confinement values starting at the same initial position. Resulting equilibrium positions are compared to results obtained by Lashgari *et al.* [273]. Blue, orange, and green circles visualise the particle shape for different particle sizes.

**Table 3:** Parameters of example case 1: migration of a single rigid particle in a square duct. See Fig. 13 for an illustration of the set-up. Grid size Δ*x* and time step Δ*t* are set to 1 in simulation units.

| Parameter | Value |
| --- | --- |
| $Re$ | 100 |
| $U$ | $1/30 \Delta x / \Delta t$ |
| $\nu$ | $1/30 \Delta x^2 / \Delta t$ |
| $\rho$ | 1 |
| $a$ | $5\Delta x, 10\Delta x, 14.28\Delta x$ |
| $w$ | $50\Delta x$ |
| $L$ | $300\Delta x$ |
| $y_0$ | $2/3w$ |
| $z_0$ | $1/3w$ |

As shown in Fig. 13(c) and (d), the particle migrates to a lateral equilibrium position located on the closest face centre for all investigated values of *χ*. As *χ* increases, the equilibrium position moves closer to the centre of the channel, matching the general trend observed experimentally [81]. Excellent quantitative agreement is found between our results and those of Lashgari *et al.* [273].

### 8.2 Example case 2: migration of a single soft particle in a square duct

The second example case investigates the impact of particle deformability on lateral migration. Originally proposed by Schaaf and Stark [117], Fig. 14(a) and (b) show a single soft spherical particle in a straight duct with square-cross section with width 2*w*. The confinement is *χ* = 0.3. The channel Reynolds number is set to Re = 10, following the definition Re = *U*_max_2*w/ν* where *U*_max_ is the maximum velocity in the channel. The flow is driven by a body force to reach the desired maximum velocity *U*_max_. The deformable capsules are modelled using the neo-Hookean model in Section 5.3. Particle deformability is characterised by the Laplace number, La, which is the ratio of particle Reynolds number, Re_p_, and capillary number, Ca, and represents the ratio of the elastic forces to the intrinsic viscous force scale:

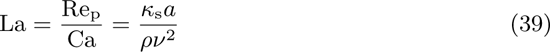

**Figure 14:**
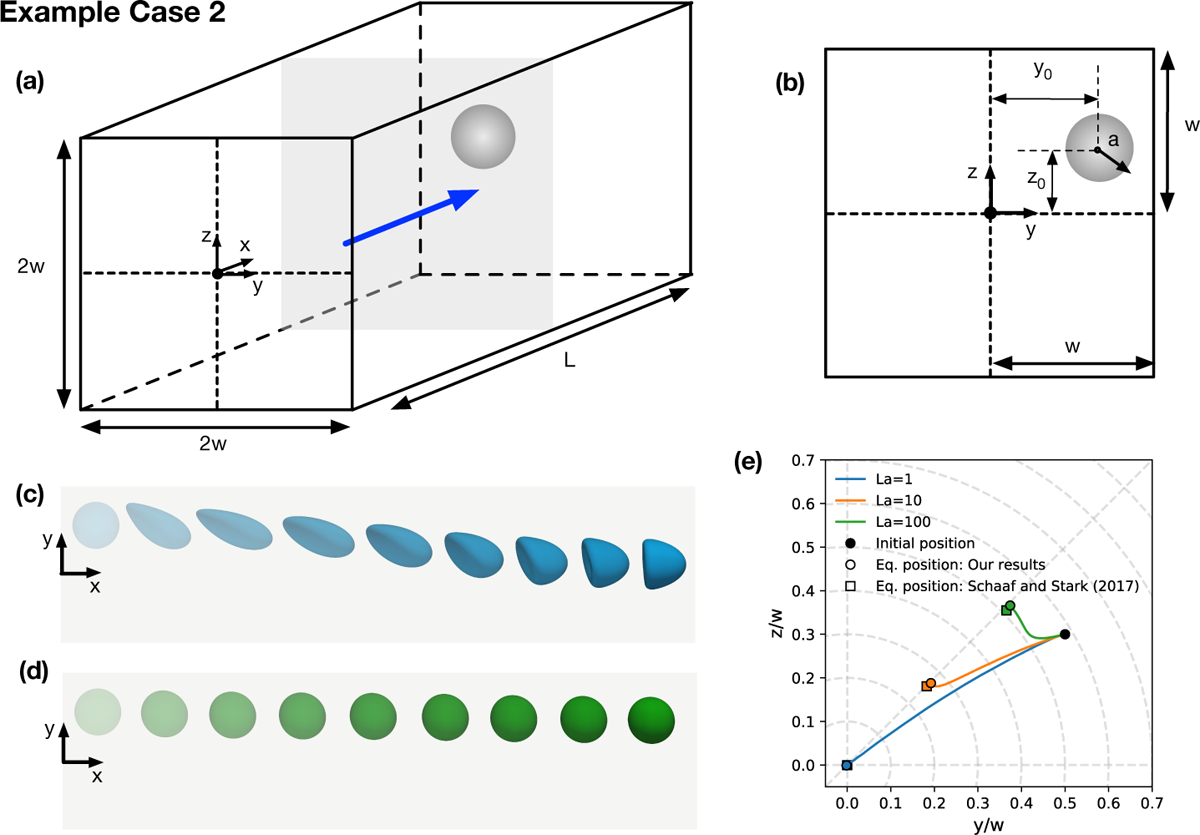
Example case 2: migration of a single soft particle in a square duct. (a) 3D schematic. (b) 2D cross-sectional schematic. (c) Migration path of a particle with La = 1 and (d) La = 100; multiple time instances are overlaid with higher saturation indicating later time. Note that the axial direction is not to scale. (e) Lateral migration paths for different Laplace numbers starting at the same initial position. Resulting equilibrium positions are compared to results obtained by Schaaf and Stark [117].

The initial position is the same for all cases as shown in Fig. 14(b). Simulation parameters are reported in Table 4.

**Table 4:** Parameters of example case 2: migration of a single soft particle in a square duct. See Fig. 14 for an illustration of the set-up. The Laplace number is controlled by the shear elasticity via (Eq. 39). Grid size Δ*x* and time step Δ*t* are set to 1 in simulation units.

| Parameter | Value |
| --- | --- |
| $Re$ | 10 |
| $La$ | 1, 10, 100 |
| $U_{\max}$ | $0.027778 \Delta x / \Delta t$ |
| $\nu$ | $1/6 \Delta x^2 / \Delta t$ |
| $\rho$ | 1 |
| $\kappa_\alpha$ | $2 \Delta x^3 / \Delta t^2$ |
| $\kappa_b$ | $0.00287 \Delta x^5 / \Delta t^2$ |
| $a$ | $18 \Delta x$ |
| $w$ | $60 \Delta x$ |
| $L$ | $480 \Delta x$ |
| $y_0$ | $0.5w$ |
| $z_0$ | $0.3w$ |

The lateral migration path of particles with La = 1, 10, and 100 are shown in Fig. 14(c). Particles begin their migration toward their equilibrium position located on the cross-sectional diagonals. The equilibrium position is La-dependent with more deformable particles migrating to positions closer to the channel centre. This observation is in line with the findings of other studies [102, 113, 150]. Excellent quantitative agreement is found between our results and those of Schaaf and Stark [117]. Fig. 14(d) and (e) show snapshots of particles with La = 1 and La = 100, respectively.

### 8.3 Example case 3: a pair of soft particles in a shear flow

Having established the lateral migration behaviour of single rigid and soft particles, we now explore particle-particle interaction in inertial flows. Originally proposed by Doddi and Bagchi [337], the third example case consists of two deformable capsules in a shear flow as shown in Fig. 15a. The initial positions of the particles have a small offset around the centre of the shearing plane ensuring that the particles migrate toward each other. The effect of inertia is investigated by increasing the particle Reynolds number

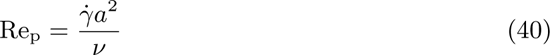

where the fluid shear rate is defined by the velocity of the confining walls (*u*_w_) and their distance *L*:

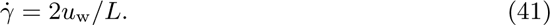

The deformable capsules are modelled using the neo-Hookean model in Section 5.3. The capillary number in (Eq. 7) characterises the capsule deformation. Simulation parameters are reported in Table 5.

**Table 5:** Parameters of example case 3: a pair of soft particles in a shear flow.

| Parameter | Value |
| --- | --- |
| $\text{Re}_p$ | 0.125, 0.575, 0.75 |
| $\text{Ca}$ | 0.025 |
| $u_w$ | 0.0090423, 0.041594, 0.054253 $\Delta x/\Delta t$ |
| $\nu$ | $1/6\Delta x^2/\Delta t$ |
| $\rho$ | 1 |
| $\kappa_b$ | $0.00287\Delta x^5/\Delta t^2$ |
| $a$ | $14.4\Delta x$ |
| $L$ | $180\Delta x$ |
| $\delta x_0$ | $8.0a = 115.2\Delta x$ |
| $\delta z_0$ | $0.4a = 5.76\Delta x$ |

As Re_p_ increases, each capsule moves closer to the mid-plane between the walls, resulting in the transition from passing to reversing trajectories. Using our in-house code, we observed this transition between Re_p_ = 0.575 and 0.75, in agreement with the results of Doddi and Bagchi [337]. Excellent agreement between the two sets of results is seen throughout the entire capsule trajectories at all values of Re_p_.

### 8.4 Example case 4: formation of a linear train of soft particles in a square duct

The final example case investigates the formation of a train of soft particles. We use the same simulation parameters as for case 2 (section 8.2) with the only difference that five particles are included in the simulation. All particles are positioned on the same side of the channel as shown in Fig. 16a. The particles are placed near the same streamline with a random variation in their lateral coordinates (*y*- and *z*-coordinates) to ensure that no ordered train exists at the beginning. The initial positions of the particles are reported in Table 6.

**Table 6:** Initial particle positions of example case 4: formation of a linear train of soft particles in a square duct. For the *x*-axis, the relative positions of each particle with respect to the first particle are provided. Absolute positions in the *y*- and *z*-directions are given with the origin on the channel centerline.

| Particle | $\delta x/a$ | $y_0/w$ | $z_0/w$ |
| --- | --- | --- | --- |
| 1 | 0 | 0.400 | 0.583 |
| 2 | -4 | 0.317 | 0.400 |
| 3 | -8 | 0.333 | 0.433 |
| 4 | -12 | 0.300 | 0.500 |
| 5 | -16 | 0.217 | 0.550 |

Fig. 16b–e show particle configurations at various times during the train formation. The time evolution of relevant spatial observables are shown in Fig. 16f–i. In the early stages, between *t/t*_ad_ = 0 and 150, the particle disorder increases significantly (Fig. 16h). During this period, multiple close particle-particle interactions lead to the swapping of lateral positions or particles passing each other. After the initial increase of disorder, particles start to band together whereby several particles follow the same general migration path, albeit with irregular fluctuations around the general trend. Once all five particles exist within a narrow lateral band, the fluctuations dampen further. Eventually, the particles migrate to their lateral equilibrium positions and form a linear train with equal inter-particle spacing throughout (Fig. 16i). One disadvantage of implementing periodic boundary conditions is demonstrated through the comparison of the absolute axial distance travelled by each particle (Fig. 16f) and the axial behaviour within the computational domain (Fig. 16i). The absolute axial distance travelled by each particle varies by several computational domain lengths (shown more clearly in the zoomed section in Fig. 16g), meaning that any given particle is interacting with periodic images upstream or downstream of it. However, valuable steady-state behaviour can still be obtained, for instance, the lateral equilibrium positions in Fig. 16h and the inter-particle spacing in Fig. 16i.

See Fig. 15 for an illustration of the set-up. The shear rate *γ* depends on Re_p_ according to (Eq. 40), and the shear elasticity *κ*_s_ is obtained from (Eq. 7). Grid size Δ*x* and time step Δ*t* are set to 1 in simulation units.

**Figure 15:**
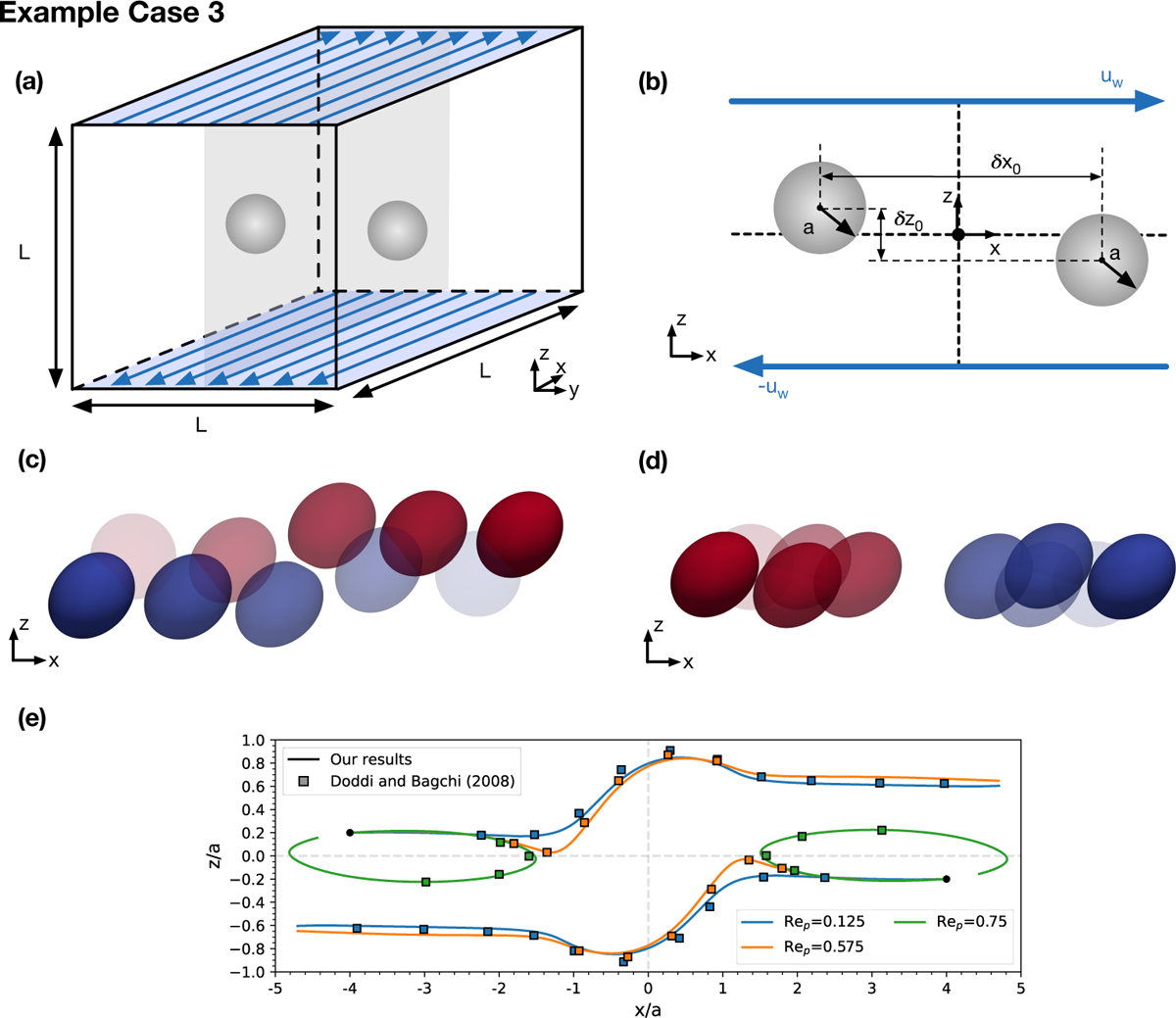
Example case 3: a pair of soft particles in a shear flow. (a) 3D schematic. (b) 2D cross-sectional schematic. Example migration paths of capsule pairs with (c) passing trajectory and (d) reversing trajectory; multiple time instances are overlaid with higher saturation indicating later time. (e) Migration paths of capsule pairs with increasing Re_p_ compared to results obtained by Doddi and Bagchi [337].

**Figure 16:**
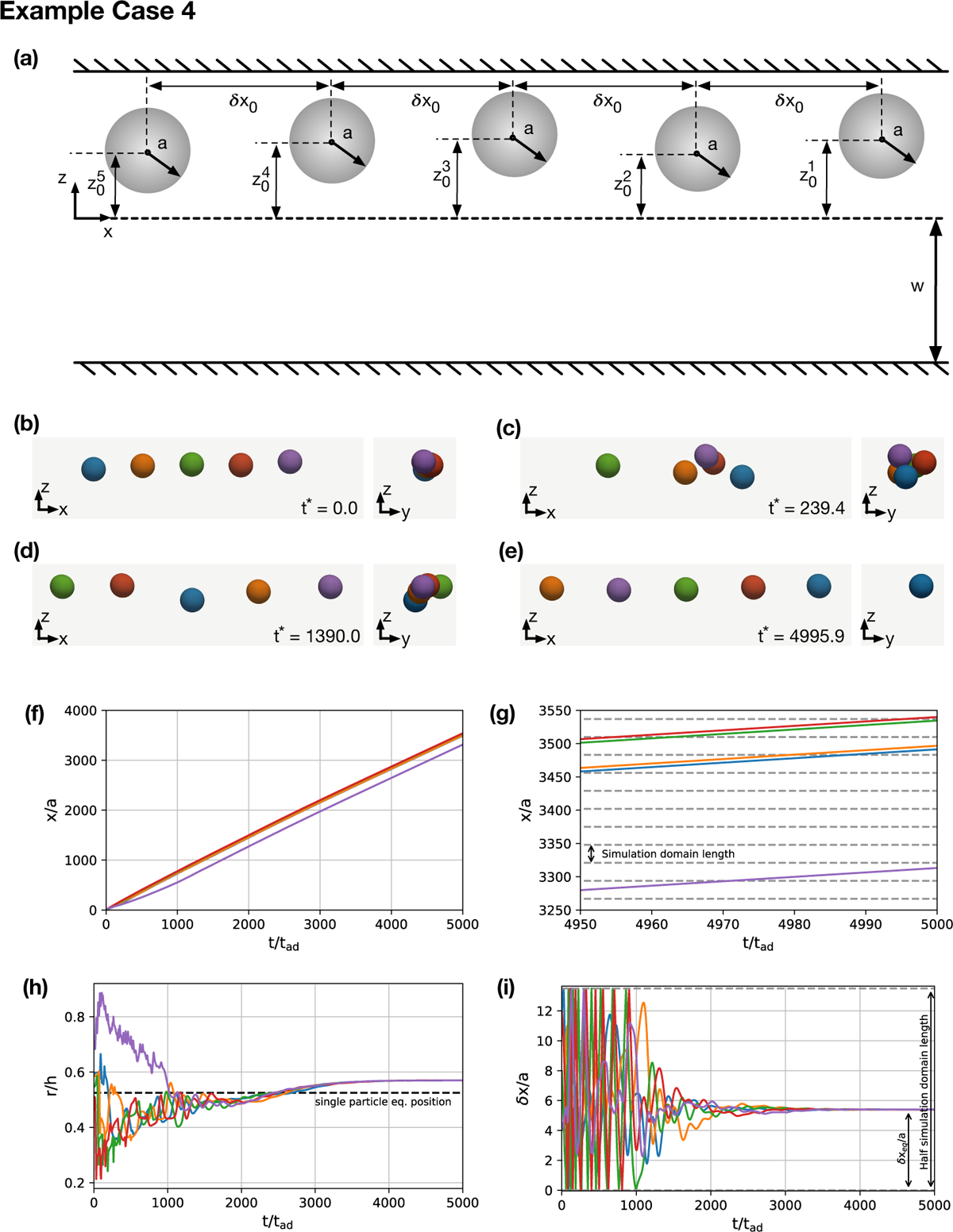
Example case 4: formation of a linear train of soft particles in a square duct. (a) 2D schematic. (b–e) Particle positions in the *x*-*z* (axial) and *y*-*z* (cross-sectional) planes at selected times. (f) Absolute axial position of each particle in time. (g) Zoomed section of absolute axial position of each particle in time. (h) Distance of each particle from the channel centreline, *r*, in time. Note that migration occurs along both the *y*- and *z*-directions. (i) Inter-particle distances within the simulation domain in time. Line colour in (f–i) corresponds to particle colour in (b–e). Note line colour in (i) refers to the leading particle in the pair.

## 9 Conclusions and Outlook

Inertial particle microfluidics (IPMF) is an emerging technology for the manipulation and separation of microparticles and biological cells. The lattice-Boltzmann (LB) method is a relatively new alternative to conventional Navier-Stokes solvers and has shown its advantages in simulating particle-laden and inertial flows. This tutorial review provides a comprehensive, yet concise overview of LB modelling for IPMF applications.

We have not attempted to replicate earlier reviews of IPMF and its applications or the LB method and its theoretical basis. Instead, we have structured this review as a top-level guide for researchers who want to employ the LB method to simulate IPMF problems. Throughout the review, we refer the reader to relevant publications for more detailed reading. We start by revisiting relevant LB-based works in terms of geometries considered (straight channels, channels with feature modifications, and curved channels) and particle concentration used (single particles, pairs and trains of particles, and non-dilute suspensions). We then describe the physical and mathematical models underpinning IPMF, including the fluid dynamics, dynamics of rigid and soft particles, boundary conditions, and relevant aspects of fluid-structure interaction (FSI). We concisely summarise the relevant numerical methods, including the LB method applied to IPMF, commonly used algorithms for the dynamics of rigid and soft particles, numerical boundary conditions, and FSI algorithms, including bounce-back variants, the momentum exchange method, and suitable immersed-boundary methods. Additionally, we provide an overview of other important simulation aspects, such as particle interaction forces, simulation initialisation, code parallelisation, grid refinement, and the selection of simulation parameters. Finally, we include four example cases that are suitable for the verification and validation of codes aiming at simulating IPMF.

Despite recent progress in the field of LB-based modelling of IPMF applications, there are several challenges and related opportunities. A key challenge is the multi-scale nature of IPMF problems. Due to the long range of inertial effects in IPMF and the need for finely resolved flow features around suspended particles, the relevant length scales range from (sub-)micron to hundreds and thousands of micrometres. Furthermore, realistic IPMF applications call for three-dimensional simulations, although two-dimensional simulations can guide the understanding of underlying effects. High-accuracy IPMF simulations in realistic geometries are, therefore, extremely computationally expensive, even with state-of-the-art parallelisation techniques. The implementation of advanced schemes for local and dynamic grid refinement and the development of well-tested reduced-order models would help overcome this challenge.

Nearly all realistic IPMF applications involve curved channels or channels with additional geometric complexity in order to generate secondary flows that accelerate the manipulation of the dynamics of suspended particles. Currently, most available algorithms and simulation codes are not suitable for this geometric complexity. A related problem is the importance of inflow and outflow boundary conditions which are essential whenever segments of an IPMF device, rather than the entire device, are simulated. To simulate the particle dynamics in a given device segment, the upstream effects need to be taken into account. Nearly all existing works ignore the history of particles accumulated in upstream segments. Moving from the commonly used flow-wise periodic boundary conditions to more realistic geometric configurations would increase the scientific value of IPMF simulations.

We also believe that the currently available methods are underused in determining the fundamental flow physics of IPMF. The vast majority of works focus on the particle kinetics, without attempting a more detailed analysis of the underlying fluid dynamics and the interaction of the particles and the fluid. Analysing IPMF problems as actually coupled fluid-particle systems would not only answer fundamental questions but also provide the physical insight needed to develop reduced-order models for less computationally demanding simulations.

We hope that this tutorial review will act as a point of entry and accompanying guide for researchers interested in LB-based modelling of IPMF problems.

## Acknowledgements

T.K. thanks Hamed Haddadi for stimulating discussions. H.S. thanks Kuntal Patel, Christopher Prohm, and Felix Rühle for collaborating on this subject.

## Author contributions

Conceptualisation: B.O. (support), S.R.B. (equal), R.V. (support), M.E.W. (equal) and T.K. (equal). Data analysis: B.O. (lead) and T.K. (support). Project administration: B.O. (equal) and T.K. (equal). Supervision: B.O. (support) and T.K. (lead). Visualisation: B.O. (equal), S.R.B. (equal), E.E. (support), Q.Z. (support) and T.K. (equal). Writing (original): B.O. (lead), K.K. (lead), S.R.B. (support), R.E. (equal), E.E. (equal), C.M. (support), F.M. (support), C.S. (equal), K.T. (support), R.V. (equal), Q.Z. (equal), M.E.W. (support), H.S. (equal) and T.K. (lead). All authors read and approved the final manuscript.

## Nomenclature

### Latin Letters

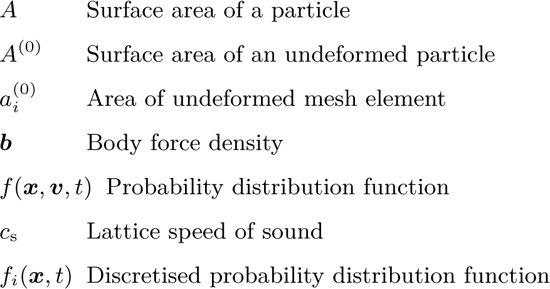
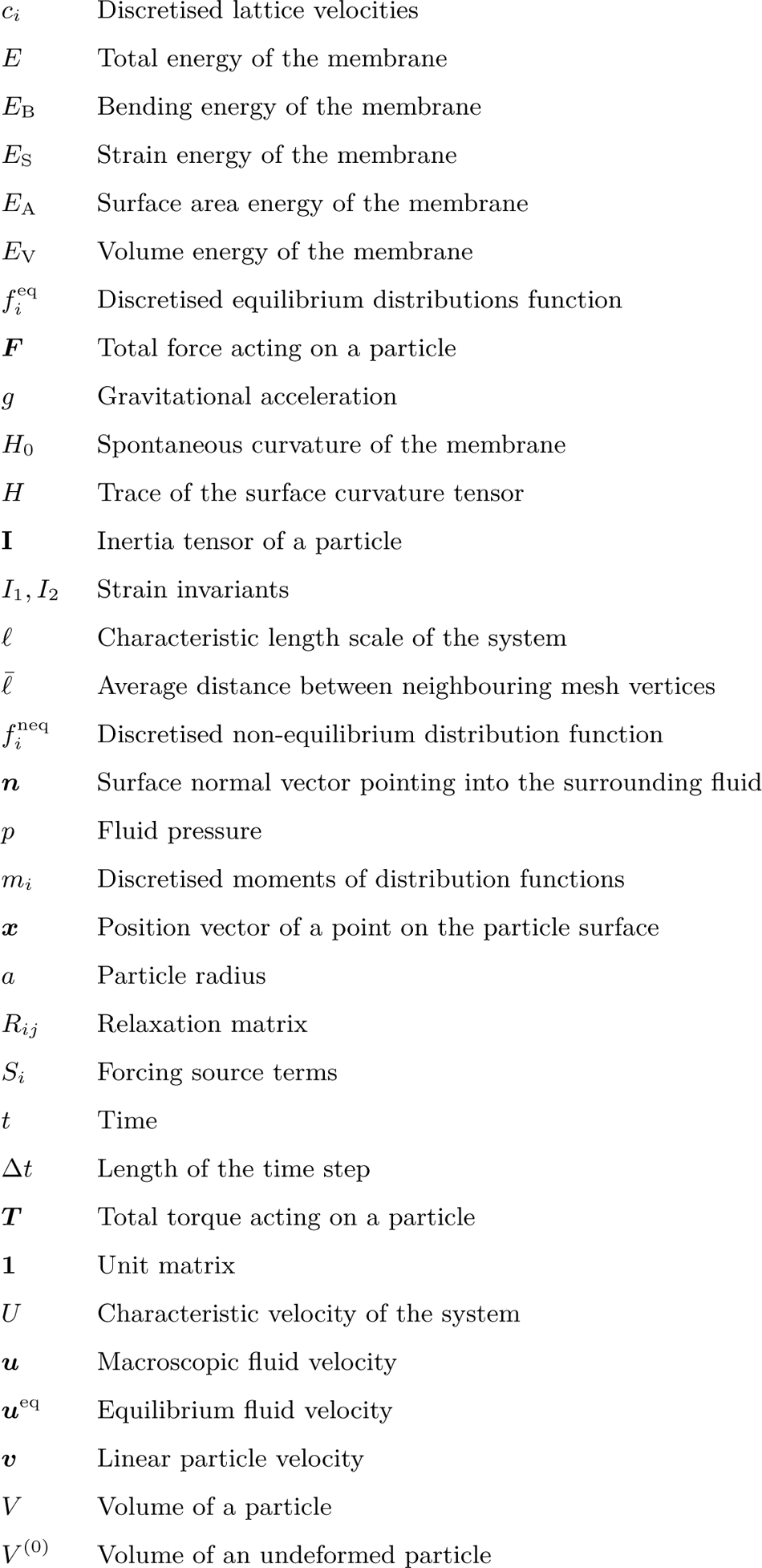

|  |  |
| --- | --- |
| $A$ | Surface area of a particle |
| $A^{(0)}$ | Surface area of an undeformed particle |
| $a_i^{(0)}$ | Area of undeformed mesh element |
| $\mathbf{b}$ | Body force density |
| $f(\mathbf{x}, \mathbf{v}, t)$ | Probability distribution function |
| $c_s$ | Lattice speed of sound |
| $f_i(\mathbf{x}, t)$ | Discretised probability distribution function |
| $c_i$ | Discretised lattice velocities |
| $E$ | Total energy of the membrane |
| $E_B$ | Bending energy of the membrane |
| $E_S$ | Strain energy of the membrane |
| $E_A$ | Surface area energy of the membrane |
| $E_V$ | Volume energy of the membrane |
| $f_i^{\text{eq}}$ | Discretised equilibrium distributions function |
| $\mathbf{F}$ | Total force acting on a particle |
| $g$ | Gravitational acceleration |
| $H_0$ | Spontaneous curvature of the membrane |
| $H$ | Trace of the surface curvature tensor |
| $\mathbf{I}$ | Inertia tensor of a particle |
| $I_1, I_2$ | Strain invariants |
| $\ell$ | Characteristic length scale of the system |
| $\bar{\ell}$ | Average distance between neighbouring mesh vertices |
| $f_i^{\text{neq}}$ | Discretised non-equilibrium distribution function |
| $\mathbf{n}$ | Surface normal vector pointing into the surrounding fluid |
| $p$ | Fluid pressure |
| $m_i$ | Discretised moments of distribution functions |
| $\mathbf{x}$ | Position vector of a point on the particle surface |
| $a$ | Particle radius |
| $R_{ij}$ | Relaxation matrix |
| $S_i$ | Forcing source terms |
| $t$ | Time |
| $\Delta t$ | Length of the time step |
| $\mathbf{T}$ | Total torque acting on a particle |
| $\mathbf{1}$ | Unit matrix |
| $U$ | Characteristic velocity of the system |
| $\mathbf{u}$ | Macroscopic fluid velocity |
| $\mathbf{u}^{\text{eq}}$ | Equilibrium fluid velocity |
| $\mathbf{v}$ | Linear particle velocity |
| $V$ | Volume of a particle |
| $V^{(0)}$ | Volume of an undeformed particle |

#### Greek Letters

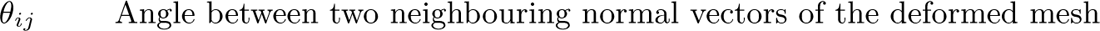
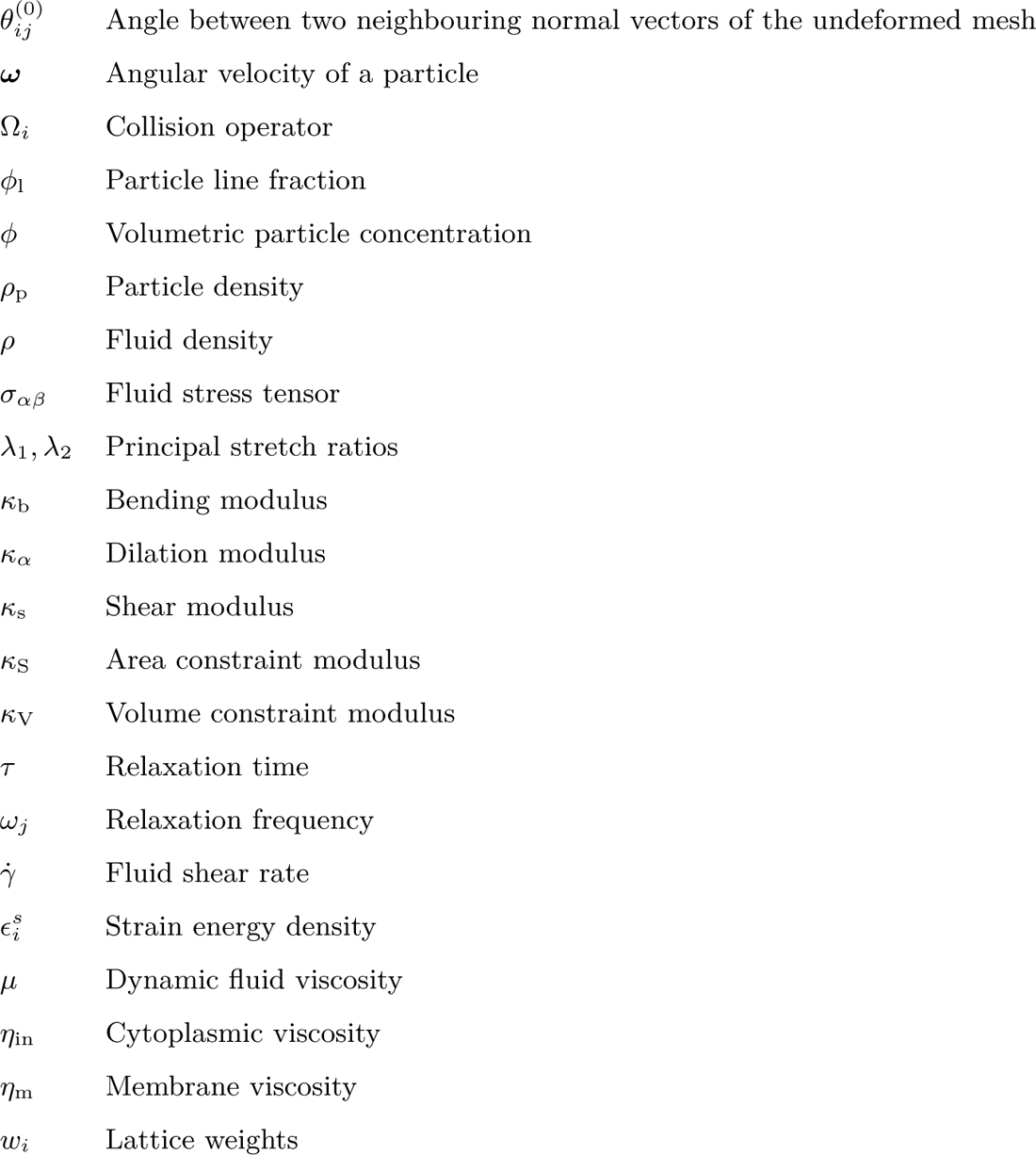

#### Superscripts

|  |  |
| --- | --- |
| Ca | Capillary number |
| $\chi$ | Particle-to-channel confinement |
| Re | Reynolds number |
| Re<sub>p</sub> | Particle Reynolds number |
| St | Stokes number |

---

**Table A1:**
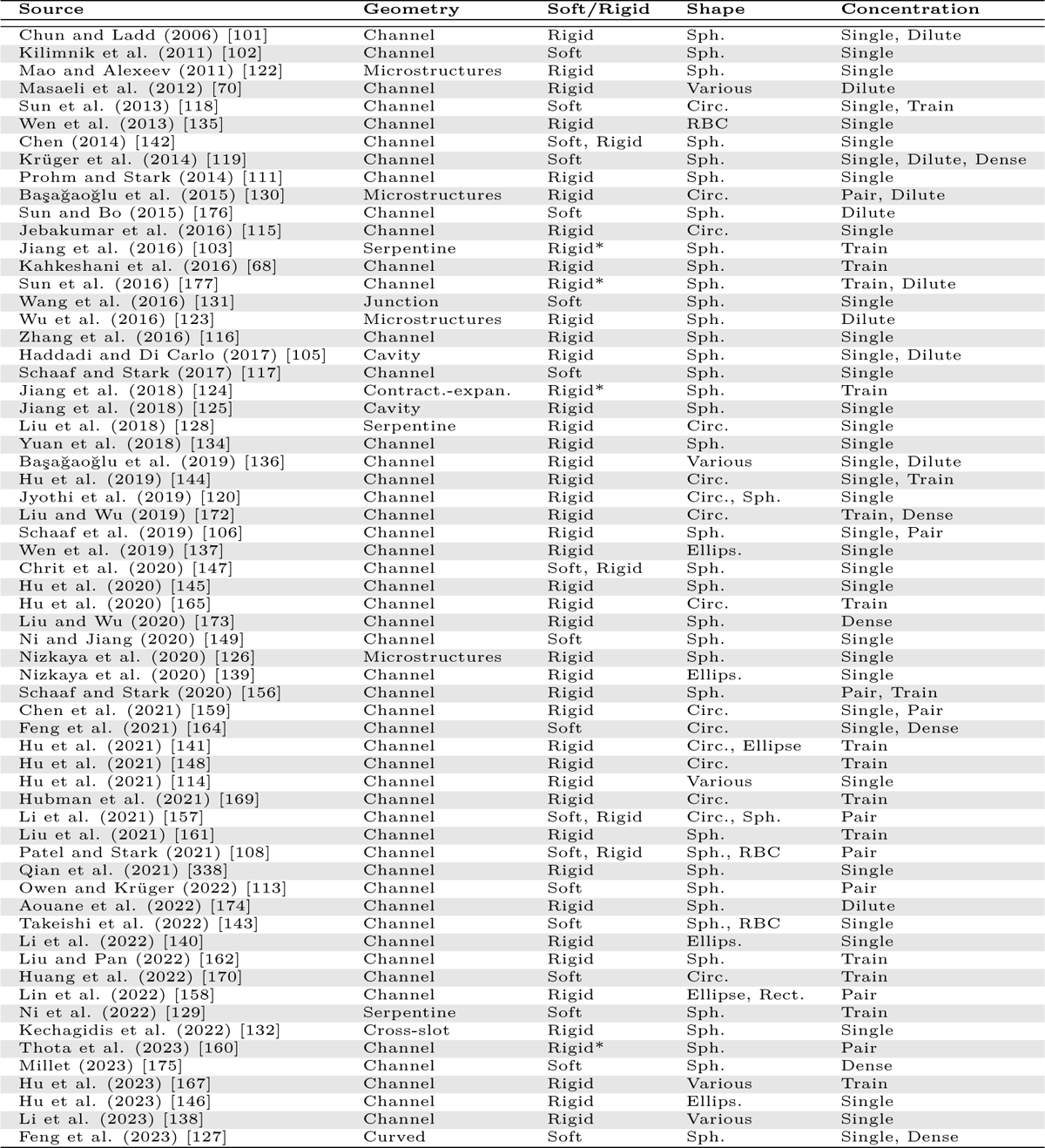
List of all LB-based works of IPMF where at least one particle is considered in a microfluidic channel. Abbreviations: Circ. = circular, Sph. = spherical, RBC = red blood cell, Ellips. = ellipsoid, Dilute = dilute suspension, Dense = dense suspension, Rigid* = rigid limit of a soft model. Note microstructures refers to any channel which features a non-smooth wall.

